# Systematic genetic mapping of necroptosis identifies SLC39A7 as modulator of death receptor trafficking

**DOI:** 10.1101/290718

**Authors:** Astrid Fauster, Manuele Rebsamen, Katharina L. Willmann, Adrian César-Razquin, Enrico Girardi, Johannes W. Bigenzahn, Fiorella Schischlik, Stefania Scorzoni, Manuela Bruckner, Justyna Konecka, Katrin Hörmann, Leonhard X. Heinz, Kaan Boztug, Giulio Superti-Furga

**Author notes:** These authors contributed equally to this work (Astrid Fauster, Manuele Rebsamen). **Corresponding Authors:** Giulio Superti-Furga CeMM Research Center for Molecular Medicine of the Austrian Academy of Sciences Lazarettgasse 14, AKH BT25.3, 1090 Vienna, Austria; Manuele Rebsamen CeMM Research Center for Molecular Medicine of the Austrian Academy of Sciences Lazarettgasse 14, AKH BT25.3, 1090 Vienna, Austria.

## Abstract

Regulation of cell and tissue homeostasis by programmed cell death is a fundamental process with wide physiological and pathological implications. The advent of scalable somatic cell genetic technologies creates the opportunity to functionally map these essential pathways, thereby identifying potential disease-relevant components. We investigated the genetic basis underlying necroptotic cell death by performing a complementary set of loss- and gain-of-function genetic screens. To this end, we established *FADD*-deficient haploid human KBM7 cells, which specifically and efficiently undergo necroptosis after a single treatment with either TNFα or the SMAC mimetic compound birinapant. A series of unbiased gene-trap screens identified key signaling mediators, such as TNFR1, RIPK1, RIPK3, and MLKL. Among the novel components, we focused on the zinc transporter SLC39A7, whose knock-out led to necroptosis resistance by affecting TNF receptor trafficking and ER homeostasis. Orthogonal, solute carrier (SLC)-focused CRISPR/*Cas9*-based genetic screens revealed the exquisite specificity of SLC39A7, among ~ 400 SLC genes, for TNFR1- and FAS-but not TRAIL-R1-mediated responses. The newly established cellular model also allowed genome-wide gain-of-function screening for genes conferring resistance to necroptosis via the CRISPR/*Cas9* synergistic activation mediator approach. Among these, we found cIAP1 and cIAP2, and characterized the role of TNIP1 (TNFAIP3-interacting protein 1), which prevented pathway activation in a ubiquitin-binding dependent manner. Altogether, the gain- and loss-of-function screens described here provide a global genetic chart of the molecular factors involved in necroptosis and death receptor signaling, prompting investigation of their individual contribution and potential role in pathological conditions.

## INTRODUCTION

Regulated cell death programs are crucial for homeostasis in multicellular organisms by eliminating cells that have become obsolete, damaged or infected(1). Necroptosis is a form of programmed necrotic cell death activated by different death and immune receptors in response to pathogen infection or in the context of sterile inflammation (2–4). Necroptotic signaling relies on activation of Receptor-interacting serine/threonine-protein kinase (RIPK)3(5–7) and its substrate Mixed lineage kinase domain-like protein (MLKL)(8, 9), which in turn mediates membrane destabilization and rupture. Death receptor ligation triggers RIPK3 activation through RIP homotypic interaction motif (RHIM) domain-mediated interaction with RIPK1, in conditions where the default apoptosis signaling pathway is blocked by caspase inhibition or absence of the adaptor protein Fas associated via death domain (FADD)(5, 10, 11). Evidence for the involvement of necroptosis in different pathologies(4, 12, 13), such as atherosclerosis(14), stroke(15) and other ischemia-reperfusion mediated injuries(16), has accumulated over the past years. While contributing to exacerbating these inflammatory conditions, necroptosis can be beneficial in other pathological contexts, particularly in restraining viral and bacterial infections(4, 17). Consequently, detailed insight into the molecular framework of necroptotic cell death entails the appealing prospect of therapeutic benefit(12, 13).

Forward genetics with somatic cells constitutes a powerful, unbiased approach to unravel the genetic basis underlying fundamental biological processes. RNA interference approaches have been successfully employed for the identification of the core necroptosis pathway members RIPK3 and MLKL(5, 6, 9). By allowing efficient generation of full knockouts through insertional mutagenesis, the near-haploid KBM7 cell line has further empowered screening approaches in human cells(18, 19). The scope of genetic screening has been broadened by the advent of CRISPR/*Cas9* technology and its adaptation to gene activation and gain-of-function screening modes(20–23), such as the development of synergistic activation mediator (SAM) libraries mediating transcriptional activation of endogenous genes(23). In this study, we combine these technologies to investigate the genetic foundation of TNFα-induced necroptosis and provide a comprehensive mapping of the molecular factors controlling necroptosis signaling. We characterize the specific contributions of the zinc transporter SLC39A7 on immune receptor function and the ubiquitin-engaging protein TNIP1 on pathway activation.

## RESULTS

### A genomically engineered KBM7 *FADD*^-^ cell line undergoes necroptosis upon treatment with TNFα or the SMAC mimetic birinapant

We set out to map the genetic requirements for necroptosis signaling in human cells, employing the haploid myeloid leukemia KBM7 cell line(18, 19). In contrast to the related HAP1 cell line that lacks RIPK3 expression(24), KBM7 are proficient at undergoing necroptosis (Figure 1a, Supplementary Figure 1a). Combined treatment with TNFα, the SMAC mimetic birinapant(25) and the pan-caspase inhibitor z-VAD-FMK induces phosphorylation of the necroptosis signaling mediators RIPK1, RIPK3, and MLKL (Supplementary Figure 1a). Pathway activation can be blocked by the RIPK1 Inhibitor II 7-Cl-ONec-1 (Nec1-s)(26) (Figure 1a, Supplementary Figure 1a). As apoptosis inhibition is required for death receptor-induced necroptosis(27), we genetically abrogated the extrinsic apoptosis pathway by deleting the signaling adaptor FADD by CRISPR/Cas9 gene editing (Supplementary Figure 1b-c). *FADD*-deficient KBM7 clones were enriched via selection for resistance to Fas ligand (FASL)- and TNF-related apoptosis-inducing ligand (TRAIL)-induced apoptosis (Supplementary Figure 1c-d). We selected a knockout subclone carrying a >100 bp insertion in the sgRNA target site, abrogating FADD expression (Supplementary Figure 1e). As expected, absence of FADD did not affect TNFα-induced activation of the canonical NF-κB pathway (Supplementary Figure 1f). Necroptosis could be induced in KBM7 *FADD*^-^ cells using TNFα or a combination of TNFα and SMAC mimetic, without requirement for caspase inhibition (Figure 1a). Indeed, cell death was blocked by Nec1-s or by MLKL inhibition using necrosulfonamide (NSA)(8), whereas apoptosis inhibition by z-VAD-FMK showed no rescue effect (Figure 1b). We further confirmed that combined treatment with TNFα and SMAC mimetic triggers rapid phosphorylation of MLKL in KBM7 *FADD*^-^ cells, indicative of necroptosis signaling, whereas it prompts apoptosis induction in KBM7 *wildtype* cells, as evidenced by Caspase-3 (CASP3) cleavage (Supplementary Figure 1g). Interestingly, we found that treatment with the SMAC mimetic birinapant alone sufficed to induce necroptosis in KBM7 *FADD*^-^ cells (Figure 1a). In the following, these newly established *FADD*-deficient KMB7 cells were used to interrogate the genetic basis of necroptotic cell death by employing complementary forward genetics approaches (Figure 1c).

**Figure 1:**
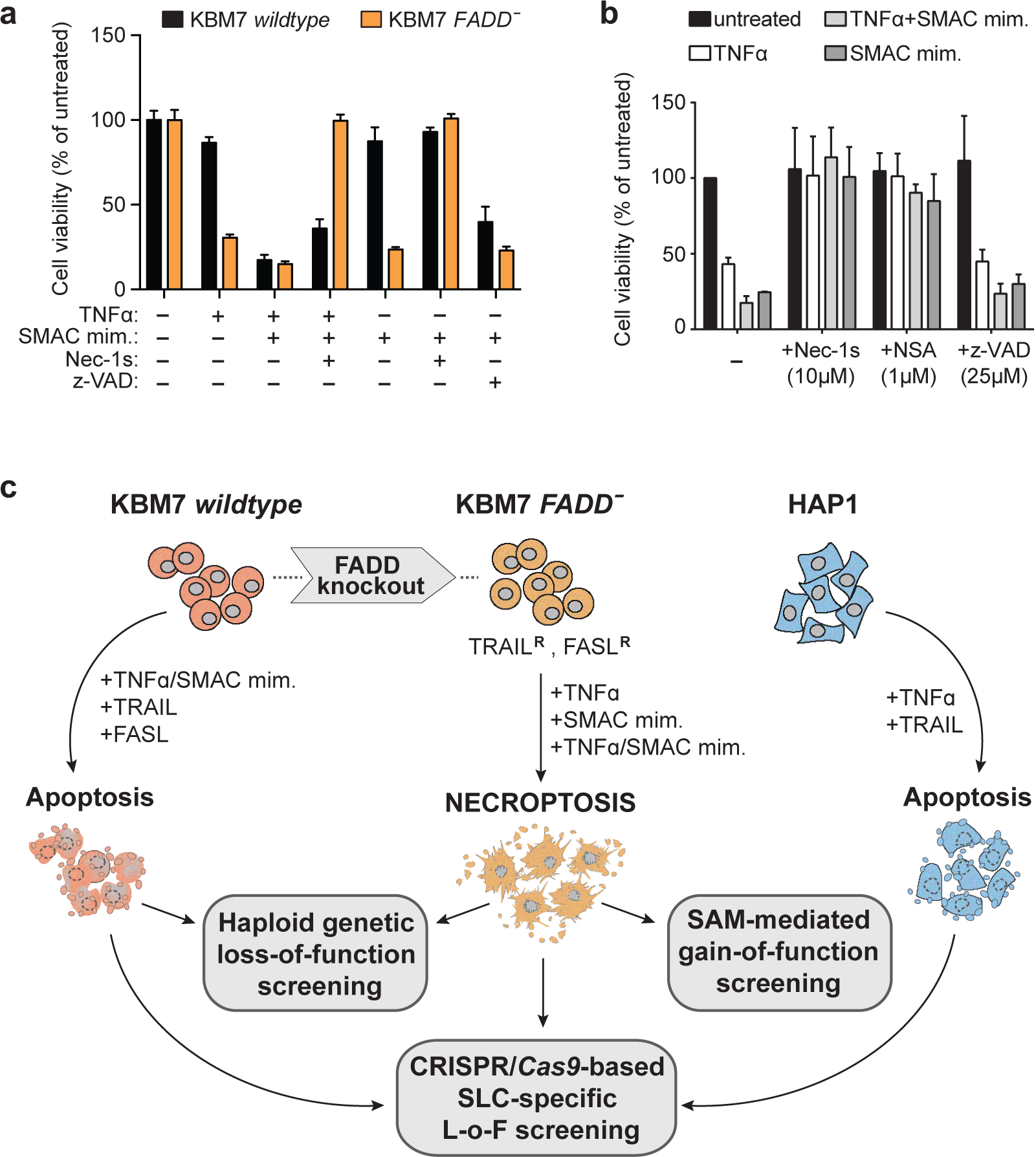
KBM7 *FADD^-^* cells undergo necroptosis upon treatment with TNFα or the SMAC mimetic birinapant. (**a**) Cell viability of KMB7 *wildtype* and KBM7 *FADD*^-^ cells treated for 24h with 100 ng/ml TNFα, 1 µM SMAC mimetic birinapant, 50 µM Nec-1s and 10 µM z-VAD-FMK as indicated. (**b**) Cell viability of KBM7 *FADD*^-^ cells treated for 24h with 100 ng/ml TNFα, 10 µM SMAC mimetic or 100 ng/ml TNFα and 1 µM SMAC mimetic, and the indicated inhibitors. Cell viability was assessed using a luminescence-based readout for ATP (CellTiter Glo) and normalized to untreated control. Data represent mean value ± s.d. of two independent experiments performed in triplicates. (**c**) Schematic overview summarizing the genetic screens presented in this study.

### Haploid genetic screens in KBM7 *FADD*^-^ cells identify the requirements for necroptosis

For identification of genes required for necroptosis signaling by haploid genetic screening, KBM7 *FADD-*cells were subjected to insertional mutagenesis with a retroviral gene-trap vector(18, 19). The insertions in the unselected mutagenized cells were mapped by deep sequencing and served as reference data set. To select for insertions conferring resistance to necroptosis, we exposed mutagenized KBM7 *FADD*^-^ cells to a high dose of the SMAC mimetic birinapant, TNFα, or a combination thereof. Each of these screens resulted in significant (*p*-value <10^-10^) enrichment of disruptive insertions in 10-13 different genes (Figures 2a-c and Supplementary Table 1). The screens identified *TNFRSF1A*, *RIPK1*, *RIPK3*, and *MLKL* among the top hits (Figure 2d) with a high number of independent insertions (Supplementary Figure 2a), consistent with their well-established role in TNFα-induced necroptosis signaling (Supplementary Figure 2b). Intriguingly, the Zinc transporter SLC39A7 was enriched as top hit along with these known necroptosis effector proteins (Supplementary Figure 2c). Other genes showing significant enrichment for mutagenic insertions were the transcription factors Wilms tumor protein (*WT1*), ETS-related transcription factor Elf-1 (*ELF1*) and Elf-4 (*ELF4*), Sp1 (*SP1*), Sp3 (*SP3*), and PU.1 (*SPI1*), as well as the Poly(rC)-binding protein 2 (*PCBP2*), Tumor necrosis factor receptor superfamily member 1B (*TNFRSF1B*), GRB2-associated-binding protein 2 (*GAB2*), Ragulator complex protein LAMTOR1 (*LAMTOR1*), and transcripts *RP11-750H9.5* and *RP11-793H13.8*. Interestingly, enrichment of insertions in *TNFRSF1B* and *SP1* was observed only for cells selected with the SMAC mimetic birinapant (Figure 2d). For follow-up analyses, we thus focused on the highly or selectively enriched genes, employing a CRISPR/*Cas9*-based multi-color competition assay (MCA) co-culture system for validation (Supplementary Figure 2d). Both SMAC mimetic as well as TNFα treatment strongly selected for GFP^+^ sg*SLC39A7*-harboring cells over control mCherry^+^ cells harboring sg*Ren* (targeting *Renilla luciferase*), indicating that loss of *SLC39A7* expression conferred enhanced cell survival or outgrowth under these selective conditions (Figure 2e). As expected, no differential outgrowth was observed when mixed populations of GFP^+^ and mCherry^+^ cells both harboring the same control sg*Ren* were analyzed. Among the further genes tested, we validated the selective advantage of cells harboring sgRNAs targeting either *SP1* or *TNFRSF1B* (Figure 2f), and, to a lesser extent, *SPI1* and *LAMTOR1* (Supplementary Figure 2e) upon treatment with the SMAC mimetic birinapant.

**Figure 2:**
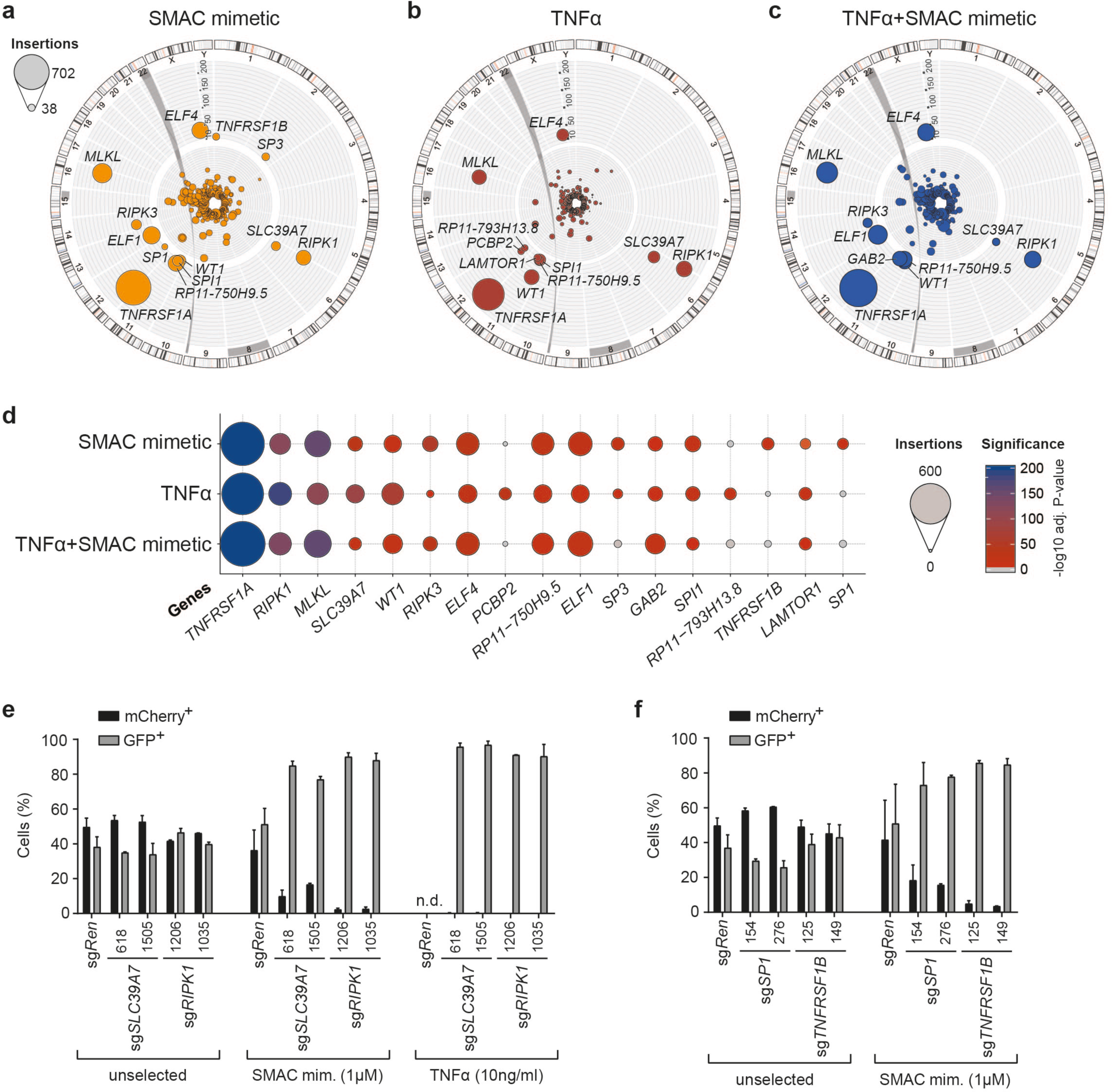
Haploid genetic screens in KBM7 *FADD*^-^ cells identify genes required for necroptosis. (**a-c**), Circos plots of haploid genetic screens in KBM7 *FADD*^-^ cells with necroptosis induction by 10 µM SMAC mimetic birinapant (**a**), 100 ng/ml TNFα (**b**), and 1 µM SMAC mimetic and 100 ng/ml TNFα combined (**c**). Each dot represents a mutagenized gene identified in the resistant cell population, dot size corresponds to the number of independent insertions identified for each gene and distance from center indicates the significance of enrichment compared to an unselected control data set. Hits with an adjusted *p*-value <10^-10^ are labelled. (**d**) Bubble plot depicting the top hits over all three screens ranked according to adjusted *p*-value. Bubble size corresponds to the number of independent insertions identified and colour gradient reflects the significance of enrichment. (**e-f**), Multi-colour competition assay (MCA) of KBM7 *FADD^-^ SpCas9* cells transduced with a GFP marker (GFP^+^) and sgRNAs targeting either *SLC39A7* or *RIPK1* (**e**), *SP1* or *TNFR2* (**f**), or *Renilla luciferase* (*sgRen*) as control, against cells transduced with sg*Ren* and an mCherry marker (mCherry^+^). The cell populations were mixed at 1:1 ratio, treated with SMAC mimetic (1 µM) or TNFα (10 ng/ml), and analyzed after 14 days by flow cytometry. Data represent mean value ± s.d. of two independent experiments performed in duplicates, n.d. (not determined) indicates wells with no outgrowth.

### Loss of SLC39A7 mediates resistance to TNFα-induced cell death by diminishing TNFR1 surface expression

We set out to investigate how loss of SLC39A7 impacts on TNFα signaling, given that it scored among the most significant hits in all the screens and that the proposed roles for this ER-resident zinc transporter did not readily explain its mechanistic link to the necroptosis phenotype(28–32). We isolated a KBM7 *FADD*^-^ subclone carrying a 5 bp deletion in the *SLC39A7* coding sequence, leading to a premature stop codon and loss of protein expression (Supplementary Figure 3a). *SLC39A7* has been reported as an essential gene in a number of cell lines of different tissue origin(33, 34), but does not form part of the core essentialome in KBM7 cells(35). Yet, loss of *SLC39A7* conferred a significant growth disadvantage as compared to sg*Ren* control cells (Figure 3a). In agreement with our screen results, *SLC39A7* knockout cells were protected from TNFα and SMAC mimetic-induced necroptosis (Figure 3b). We next investigated the impact of *SLC39A7* loss on TNFα signaling and found that knockout cells failed to activate the canonical NF-κB pathway following TNFα stimulation (Figure 3c). Strikingly, we could not detect Tumor necrosis factor receptor superfamily member 1A (TNFR1) surface expression in *SLC39A7*^-^ cells (Figure 3d). We confirmed that SLC39A7 localizes to the ER (28)(Supplementary Figure 3b–c) and, in order to assess the cellular alterations resulting from its loss, we determined changes in protein levels by employing a proteomics approach focused on membrane compartments (Supplementary Figure 3d-e). Gene ontology (GO) term enrichment among the proteins significantly upregulated in *SLC39A7*^-^ cells yielded ‘*response to endoplasmic reticulum stress*’ as top hit (Supplementary Figure 3f-g, Supplementary Table 2). Gene set enrichment analysis (GSEA) among all significantly changed targets moreover identified the hallmark gene set ‘*Unfolded_protein_response*’ (UPR) as top positive enriched hit (Supplementary Figure 3h). These data strongly indicated that, similar to its *Drosophila* and murine orthologues(30, 31, 36), loss of human SLC39A7 impacts on ER homeostasis. Indeed, *SLC39A7*^-^ cells displayed ER stress at basal state, reflected in increased levels of BiP (ER chaperone 78 kDa glucose-regulated protein GRP), and exhibited higher sensitivity towards different chemical ER stressors as monitored by induction of CHOP (DNA damage-inducible transcript 3 protein DDIT3) (Figure 3e). Given the crucial role of the ER in synthesis, folding and shuttling of membrane proteins(37), we next explored how elevated ER stress upon loss of *SLC39A7* impinged on TNFR1 trafficking. Surprisingly, while TNFR1 was not detectable on the surface of *SLC39A7*^-^ cells by flow cytometry (Figure 3d), we found higher levels of intracellular TNFR1 protein in *SLC39A7* knockouts as monitored by whole cell immunoblot analysis (Figure 3f). Using glycan maturation as a readout for glycoprotein movement through the secretory pathway(38), we found that TNFR1 accumulating in *SLC39A7*^-^ cells exhibited sensitivity to Endo H, indicative of retained ER localization (Figure 3g).

**Figure 3:**
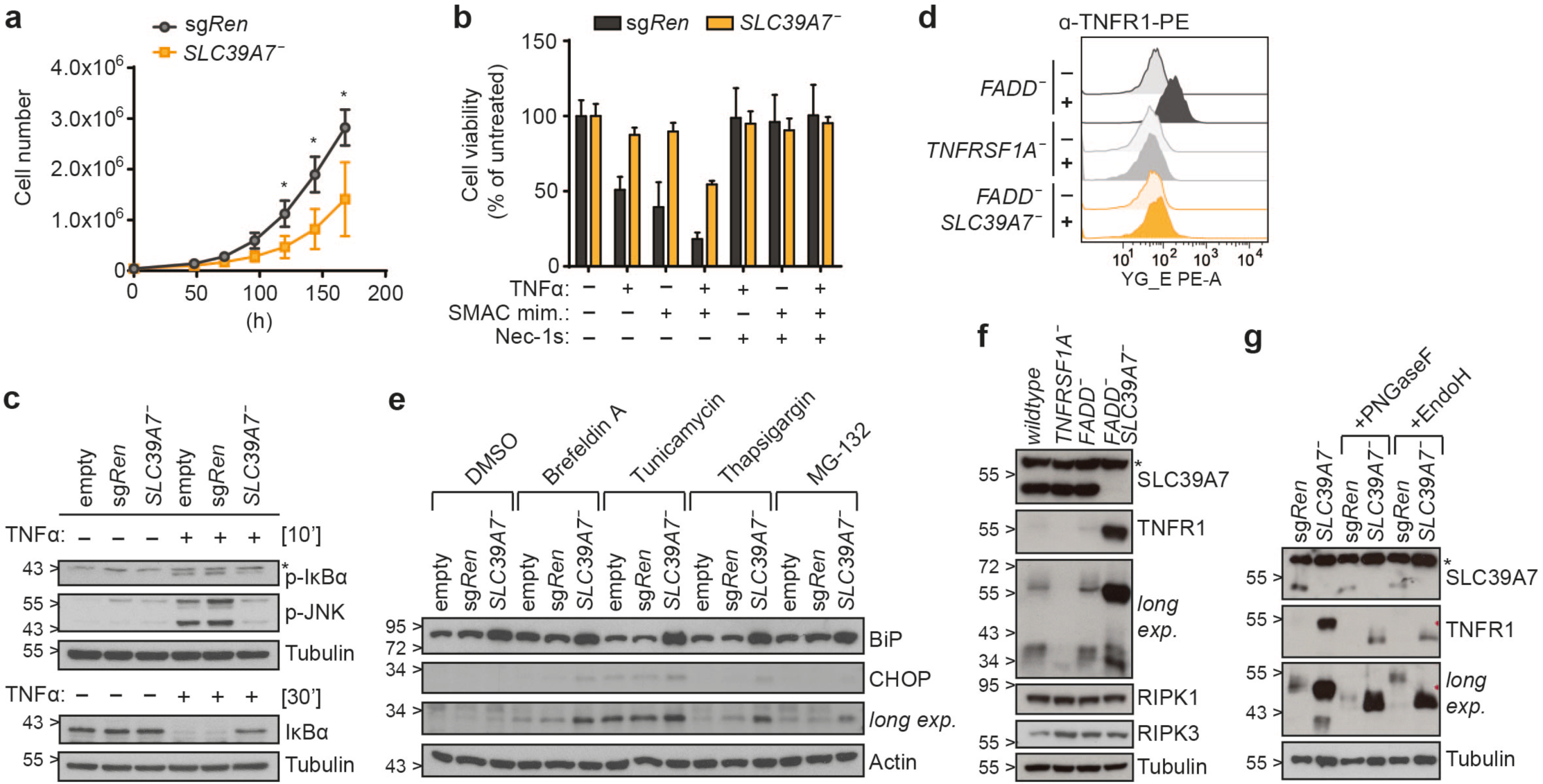
Loss of SLC39A7 mediates resistance to TNFα-induced cell death by diminishing TNFR1 surface expression. (**a**) Growth curve for KBM7 *FADD*^-^ sg*Ren* or *SLC39A7*^-^ cells. Equal cell numbers were seeded and cell growth monitored over 7 days. Data represent mean value ± s.d. of five independent experiments; statistical analysis by *t*-test, \**P*<0.01. (**b**) Cell viability in KBM7 *FADD*^-^ sg*Ren* or *SLC39A7*^-^ cells treated overnight (16h) with TNFα (10 ng/ml), SMAC mimetic (0.5 µM), and Nec-1s (50 µM) as indicated. Cell viability was assessed using a luminescence-based readout for ATP (CellTiter Glo). Data represent mean value ± s.d. of two independent experiments performed in triplicates. (**c**) KBM7 *FADD^-^ SpCas9* (empty, sg*Ren* or *SLC39A7^-^*) cells were stimulated for the time indicated with TNFα (10 ng/ml). Cells were then lysed and subjected to immunoblotting with the indicated antibodies. (**d**) Flow cytometry analysis for TNFR1 surface expression. KBM7 *TNFRSF1A*^-^ cells serve as negative control for background staining. Data shown are representative of two independent experiments. (**e**) KBM7 *FADD^-^ SpCas9* (empty, sg*Ren* or *SLC39A7^-^*) cells were treated for 7h with Brefeldin A (0.5 µM), Tunicamycin (2 µM), Thapsigargin (0.5 µM), MG-132 (10 µM) or DMSO as control. Cells were then lysed and subjected to immunoblotting with the indicated antibodies. (**f**) KBM7 *TNFRSF1A^-^*, KBM7 *FADD*^-^ and KBM7 *FADD^-^ SLC39A7*^-^ cells were lysed and subjected to immunoblotting with the indicated antibodies. (**g**) KBM7 *FADD^-^ SpCas9* sg*Ren* or *SLC39A7*^-^ cell lysates were incubated for 1h at 37°C in presence or absence of PNGaseF or EndoH, respectively, and analysed by immunoblot with the indicated antibodies. Immunoblots shown are representative of two independent experiments, * indicates non-specific band.

### Orthogonal genetic screens and surface marker analysis define the specificity of SLC39A7 on receptor trafficking

To study the specificity of the *SLC39A7* loss-of-function-induced defect in TNFR1 trafficking, we devised a set of targeted and genome-wide screens including the other prominent death domain-containing members of the TNF receptor superfamily, Fas Cell Surface Death Receptor (FAS) and TNF-Related Apoptosis-Inducing Ligand Receptor (TRAIL-R)1 and 2. The use of a focused CRISPR/*Cas9* library targeting the 388 members of the SLC family allowed us to efficiently screen various conditions and stimuli in multiple cell lines. KBM7 *FADD^-^*, KBM7 *wildtype* and HAP1 cells were lentivirally transduced with the library and subsequently selected for resistance towards cell death induction using different doses of TNFα, SMAC mimetic, TRAIL, or FASL (Supplementary Figure 4a). sgRNAs targeting SLC39A7 were top-enriched in screens with TNFα, SMAC mimetic, and, interestingly, FASL (Figure 4a, Supplementary Table 4 and 5), highlighting the exquisite role of SLC39A7 among SLCs in affecting these crucial immune signaling pathways. Intriguingly, loss of *SLC39A7* did not confer a selective advantage when TRAIL was used to induce cell death. Genome-wide haploid genetic screening in KBM7 cells confirmed the divergent requirement for SLC39A7 between TRAIL- and FASL-induced cell death (Supplementary Figure 4b, Supplementary Table 1). We validated these contrasting phenotypes by MCA in Jurkat E6.1 (Figure 4b) and KBM7 cells (Supplementary Figure 4c). In line with the screen findings, treatment with FASL selected for GFP^+^ sg*SLC39A7*-harboring cells, whereas we found no differential outgrowth in untreated or TRAIL-selected conditions. Similar to TNFR1, surface expression of FAS and TRAIL-R2 was reduced in *SLC39A7* knockout cells (Figure 4c and Supplementary Figure 4d). In contrast, TRAIL-R1 was still detectable and even slightly increased in these cells (Figure 4c). While explaining the retained sensitivity to TRAIL-induced cell death, the dichotomy between these two closely related receptors was surprising. Consequently, we monitored a wider panel of cell surface proteins present on KBM7 cells (Supplementary Figure 4e)(39). Several markers, including CD31, CD11a, CD34, CD45 and CD55 exhibited altered surface expression in *SLC39A7* knockout cells, while others, such as CD4, C3AR or C5L2, remained unchanged. Taken together, these data indicate that specific membrane proteins are differentially affected by loss of *SLC39A7*.

**Figure 4:**
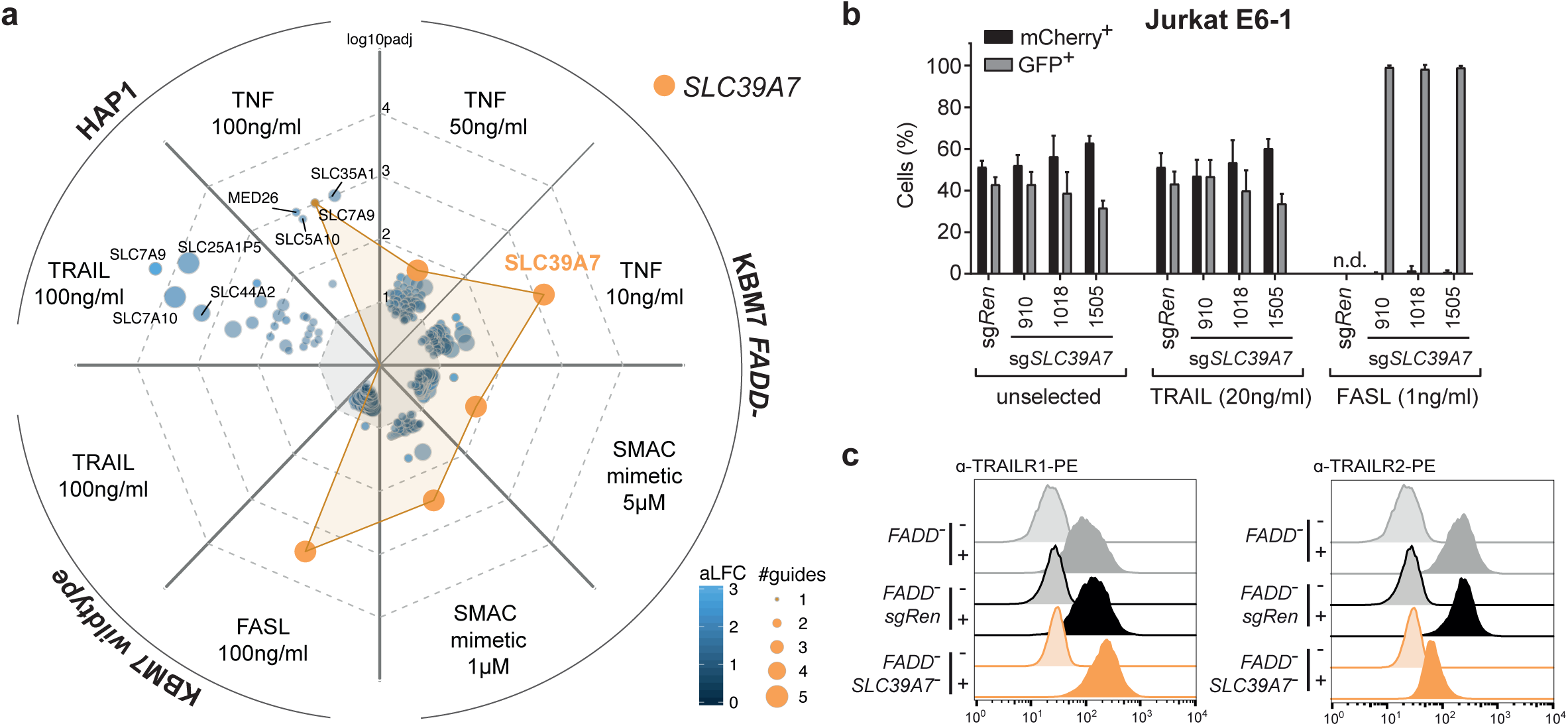
Orthogonal genetic screens and surface marker analysis specify SLC39A7 loss-of-function phenotype on receptor trafficking. (**a**) Spider plot summarizing the results of 8 independent CRISPR/*Cas9* screens in KBM7 *wildtype*, KBM7 *FADD*^-^ or HAP1 cells using an SLC-specific gRNA library. Each plot section represents one screen with the indicated stimuli. Screen analysis was performed by identifying differentially enriched sgRNAs using DESeq2 and then aggregating sgRNAs to genes using Gene Set Enrichment Analysis. Identified hits are ranked according to the adjusted *p*-value of enrichment (–log10(p_adj_)), bubble size indicates the number of significantly enriched sgRNAs and colour the average log2 fold-change (aLFC) of the enriched sgRNAs. Screens were performed in duplicate. *SLC39A7* is highlighted in orange. No gene was identified in KBM7 *wildtype* cells upon TRAIL treatment (**b**) MCA of Jurkat E6.1 *SpCas9* cells transduced with a GFP marker (GFP^+^) and sgRNAs targeting either *SLC39A7* or *Renilla luciferase*(sg*Ren*) as control, against cells transduced with sg*Ren* and an mCherry marker (mCherry^+^). The cell populations were mixed at 1:1 ratio, treated with TRAIL (20 ng/ml) or FASL (1 ng/ml), and analyzed after 14 days by flow cytometry. Data represent mean value ± s.d. of two independent experiments performed in duplicates, n.d. (not determined) indicates wells with no outgrowth. (**c**) Flow cytometry analysis for TRAILR1 (left) and TRAILR2 (right) surface expression in KBM7 *FADD^-^,* KBM7 *FADD*^-^ sg*Ren* or KBM7 *FADD^-^ SLC39A7*^-^ cells. Data shown are representative of two independent experiments.

### SLC39A7 transporter function is required for ER homeostasis, TNFR1 surface expression and cell death induction

To corroborate the link between *SLC39A7* loss and the phenotypes observed, we reconstituted KBM7 *FADD^-^ SLC39A7* knockout cells with V5-tagged SLC39A7 or GFP control (Figure 5a). SLC39A7 reconstitution decreased BiP expression, restored TNFR1 surface levels (Figure 5b) and, consequently, sensitivity to cell death induction with TNFα and SMAC mimetic (Figure 5c). To investigate the requirement of SLC39A7 transport activity, we performed analogous reconstitution experiments with SLC39A7 constructs bearing substitutions at conserved histidines (H329A or H358A) predicted to participate in an intramembranous zinc-binding site or at other conserved residues (H362A or G365R) within the same metalloprotease-like motif(40). SLC39A7 H329A and H358A mutants failed to restore sensitivity to necroptosis (Figure 5d) and to relieve ER stress (Figure 5e), whereas H362A and G365R behaved similar to the *wildtype* construct. These results are consistent with recent structural and transport data for SLC39A4 (ZIP4), in which substitution of histidines corresponding to SLC39A7 H329 and H358, but not H362, was shown to affect transport activity(41). Compared to the constructs able to functionally rescue SLC39A7 deficiency, H329A and H358A mutants showed reduced expression in KBM7 FADD^-^ SLC39A7^-^ cells (Figure 5e). This is likely a consequence of the unresolved ER stress in these cells, as all constructs express at comparable level in SLC39A7-proficient cells (Figure 5e). While an intrinsic reduced stability of H329A and H358A mutants cannot be excluded, their expression was comparable to endogenous SLC39A7 levels, strongly suggesting that transport activity is required to restore ER homeostasis and necroptosis sensitivity.

**Figure 5:**
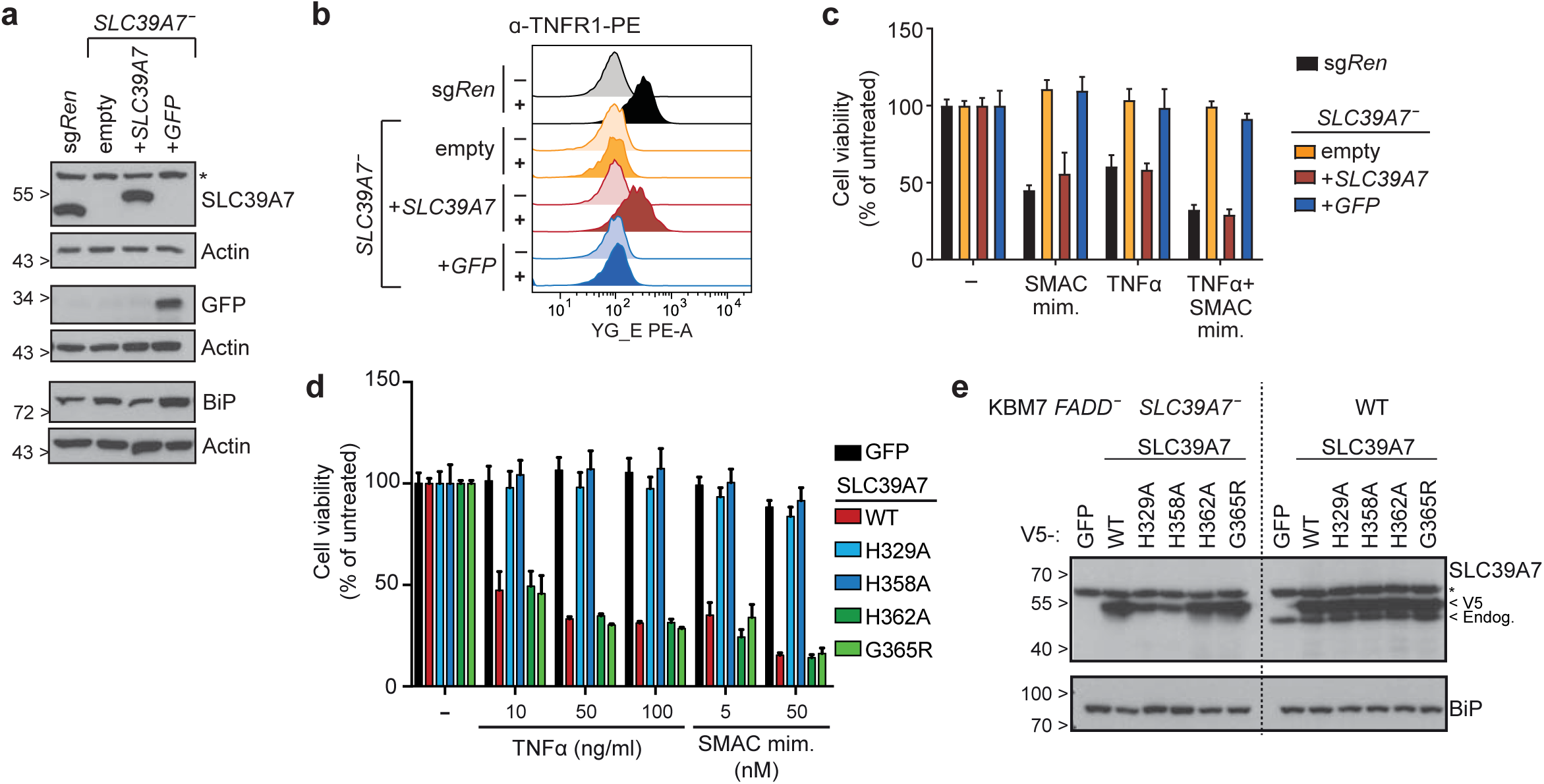
SLC39A7 transporter function is required to relieve ER stress, restore TNFR1 surface expression and induce cell death. (**a**) KBM7 *FADD^-^ SLC39A7*^-^ cells lentivirally transduced with either SLC39A7 or GFP were lysed and immunoblotted with the indicated antibodies, KBM7 *FADD*^-^ sg*Ren* cells serve as control. Immunoblots shown are representative of two independent experiments, * indicates non-specific band. (**b**) Flow cytometry analysis for TNFR1 surface expression in KBM7 *FADD^-^ SLC39A7*^-^ cells reconstituted with either SLC39A7 or GFP. KBM7 *FADD*^-^ sg*Ren* and empty KBM7 *FADD^-^ SLC39A7*^-^ cells serve as positive and negative control, respectively. Data shown are representative of two independent experiments. (**c**) Cell viability in KBM7 *FADD^-^ SLC39A7*^-^ cells reconstituted with either SLC39A7 or GFP, KBM7 *FADD*^-^ sg*Ren* and empty KBM7 *FADD^-^ SLC39A7*^-^ serve as controls. Cells were treated overnight (16h) with TNFα (10 ng/ml), SMAC mimetic (0.5 µM), or a combination thereof. Cell viability was assessed using a luminescence-based readout for ATP (CellTiter Glo). Data represent mean value ± s.d. of two independent experiments performed in triplicates. (**d**) Cell viability in KBM7 *FADD^-^ SLC39A7*^-^ cells stably reconstituted with GFP or the indicated SLC39A7 constructs. Cells were treated as indicated for 24h and cell viability was assessed as in c. Data represent mean value ± s.d. of two independent experiments performed in triplicates. (**e**) KBM7 *FADD^-^ SLC39A7*^-^ or KBM7 *FADD*^-^ cells stably reconstituted with the specified constructs were lysed and subjected to immunoblotting with the indicated antibodies. * indicates non-specific band.

### Genome-scale gain-of-function screens identify negative regulators of necroptotic cell death signaling

Combination of the newly generated KBM7 *FADD*^-^ cell necroptosis model with recently developed CRISPR/*Cas9*-based technologies mediating transcriptional activation of endogenous genes offered the opportunity to identify cellular inhibitors of necroptosis. We thus performed complementary gain-of-function screens employing the CRISPR/*Cas9*-based SAM approach(23). KBM7 *FADD*^-^ cells expressing dCas9-VP64 and MS2-p65-HSF1 were transduced with the genome-scale SAM sgRNA lentiviral library and subsequently selected with two doses of either TNFα or birinapant for 72 hours. NGS sequencing and downstream analysis allowed the identification of sgRNAs specifically enriched upon necroptosis induction, thereby revealing genes conferring resistance to necroptotic cell death when upregulated (Figure 6a-b, Supplementary Table 6 and 7). Genome-wide screens for resistance against birinapant-induced cell death revealed the SMAC mimetic target protein Cellular inhibitor of apoptosis (cIAP)2 (encoded by *BIRC3*) as top scoring gene and enriched sgRNAs targeting cIAP1 (*BIRC2*), thus confirming the validity of our approach (Figure 6b). TNFα treatment similarly revealed BIRC2 among other genes, and, notably, TNIP1 (TNFAIP3 interacting protein 1, also known as ABIN-1) as the top scoring hit (Figure 6a). Identification of TNIP1 was particularly interesting as this ubiquitin-binding protein affects multiple inflammatory pathways, including TNF receptor and Toll-like receptor signaling(42, 43). Indeed, TNIP1 was shown to negatively regulate NF-kB and MAPK activation(43, 44) as well as to inhibit TNFα-induced apoptosis(45). Moreover, single nucleotide polymorphisms (SNPs) in TNIP1 have been associated with several inflammatory diseases comprising psoriasis, systemic lupus erythematosus and systemic sclerosis(42). In line with this, mice deficient for TNIP1 or knock-in for a ubiquitin-binding-defective mutant (TNIP1[D485N]) develop autoimmunity(46–49). We first generated KBM7 *FADD*^-^ cells expressing individual SAM-sgRNAs targeting *TNIP1*, *BIRC3* or *Renilla* luciferase as control. Cells overexpressing TNIP1 or BIRC3 showed increased resistance to TNFα-or birinapant-induced necroptosis, respectively, both upon short (24h) (Supplementary Figure 5a-b) and long (72h) treatment (Figure 6c-d). The protective effect of different sgRNAs correlated with the level of overexpression achieved, with the strongest sgRNA for each gene showing partial protection to both stimuli (Figure 6e). To verify that resistance to TNFα-induced necroptosis is conferred by TNIP1 and not due to off-target effects from the sgRNAs or the SAM approach, we established KBM7 FADD^-^ cells in which TNIP1 upregulation results from ectopic cDNA expression (Figure 7b). While cells overexpressing *wildtype* TNIP1 were resistant to TNFα-induced necroptosis, expression of a construct bearing a point mutation (D472A) disrupting its ubiquitin-binding domain(47) was ineffective, demonstrating that protection requires TNIP1 ubiquitin-binding activity (Figure 7a). Cells overexpressing *wildtype* but not mutant TNIP1 showed a reduced phosphorylation of RIPK1 and MLKL after TNFα stimulation (Figure 7c), indicating that TNIP1 affects the necroptosis pathway upstream of these critical signaling events. In contrast, we did not observe any major effect of TNIP1 overexpression on TNFα-induced NF-kB or MAPK pathway activation (Figure 7d). Similar results were obtained upon sgRNA-mediated TNIP1 overexpression (Supplementary Figure 5c-d). In summary, these data unveil a role for TNIP1 in controlling TNFα-induced cell death beyond apoptosis. Regulation of necroptosis could therefore contribute to the inflammatory phenotypes observed in TNIP1-deficient and TNIP1[D485N] knock-in mice, and be connected with its association with multiple inflammatory diseases.

**Figure 6:**
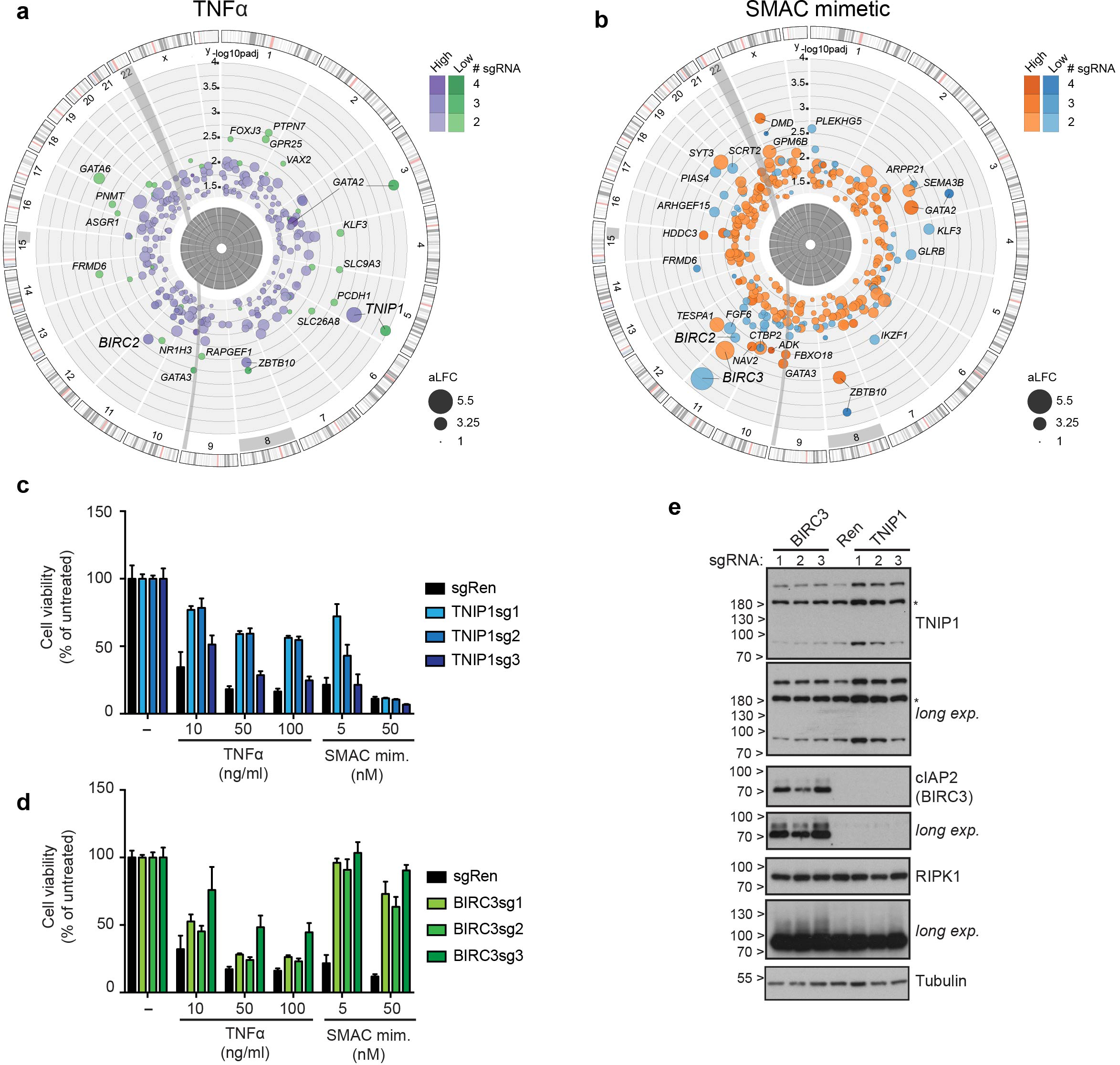
Genome-scale gene activation screens identify regulators of necroptosis. (**a**-**b**) Circos plots of genome-scale SAM screens in KBM7 *FADD*^-^ SAM cells with necroptosis induction by TNFα (**a**) or SMAC mimetic birinapant (**b**). For each stimulus, screens were performed at low and high concentrations (TNFα: 10 ng/ml (green) or 100 ng/ml (purple); birinapant: 0.1 µM (blue) or 1 µM (orange)). Screen analysis was performed by identifying differentially enriched sgRNAs using DESeq2 and then aggregating sgRNAs to genes using Gene Set Enrichment Analysis. Identified hits are ranked according to the adjusted *p*-value of enrichment (–log10(padj). Bubble size corresponds to the average log2 fold-change (aLFC) of enrichment, color indicates the number of significantly enriched sgRNAs. Screens were performed in duplicate except the high concentration of Birinipant in simplicate. **(c-d)** Cell viability in KBM7 *FADD*^-^ SAM cells transduced with sgRNA targeting *TNIP1, BIRC3* or *Renilla luciferase* (Ren). Cells were treated as indicated for 72h and viability was assessed using a luminescence-based readout for ATP (CellTiter Glo). Data represent mean value ± s.d. of two independent experiments performed in triplicates. (**e**) KBM7 *FADD^-^ CRISPR-SAM cells* transduced with the specified sgRNAs were lysed and subjected to immunoblotting with the indicated antibodies.

**Figure 7:**
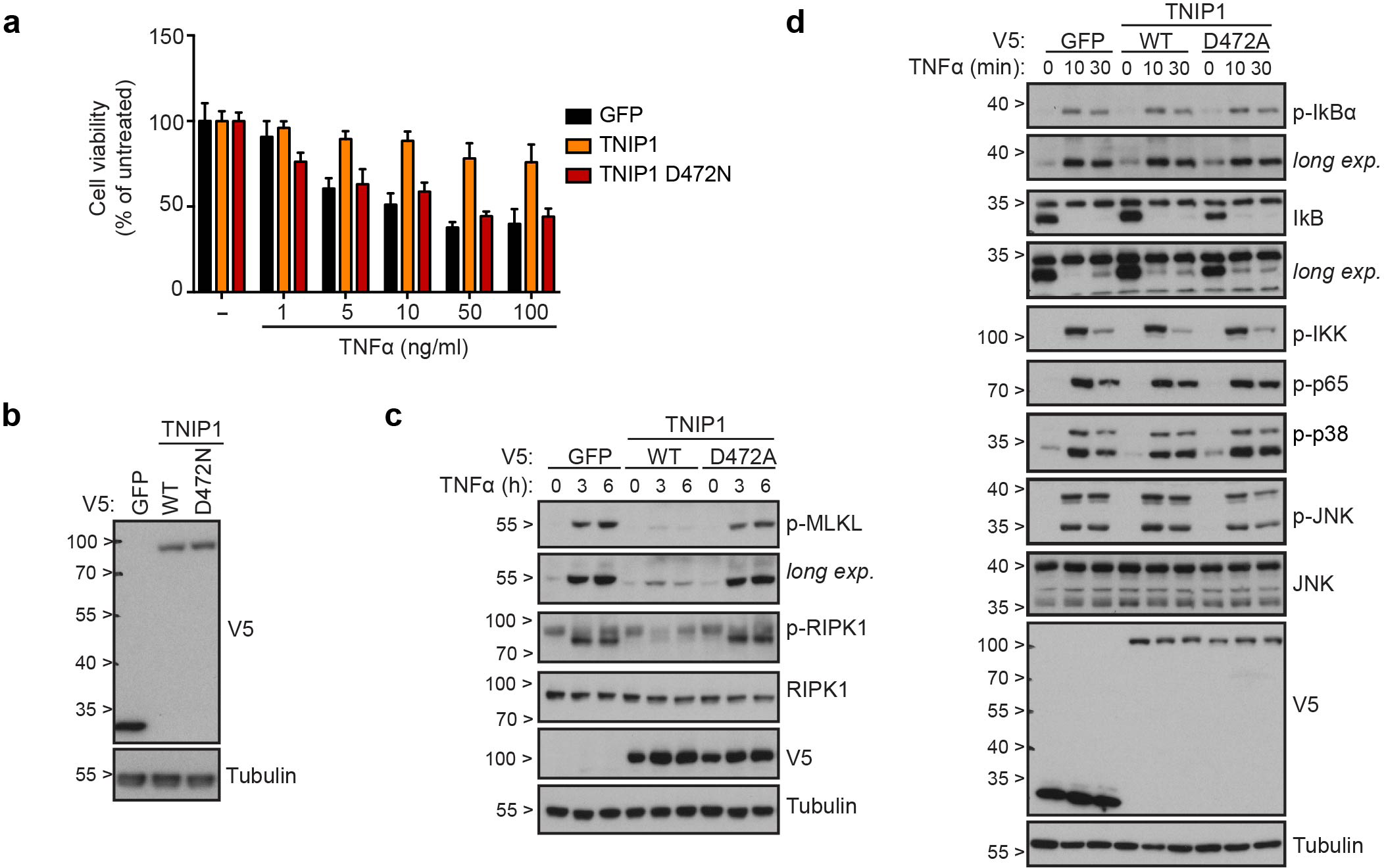
TNIP1 interferes with TNF α-induced necroptosis and RIPK1 activation in a ubiquitin binding–dependent manner. **(a-b)** Cell viability (**a**) and immunoblot (**b**) of KBM7 *FADD*^-^ cells stably expressing the specified V5-tagged TNIP1 constructs or GFP. In **a** cells were treated as indicated for 24h and viability was assessed using a luminescence-based readout for ATP (CellTiter Glo). Data represent mean value ± s.d. of two independent experiments performed in triplicates. (**c-d**) KBM7 *FADD*^-^ cells stably expressing the specified TNIP1 constructs were stimulated for the time indicated with TNFα (100 ng/ml in **c**; 10 ng/ml in **d**). Cells were then lysed and subjected to immunoblotting with the indicated antibodies. Data shown are representative of at least two independent experiments.

## DISCUSSION

Despite its relatively recent discovery, evidence for the relevance of necroptosis in human physiology and pathology has accumulated over the past decade(4, 13, 15, 50). Thus, defining the functional genetic landscape of this fundamental process is expected to expand the understanding of this pathway and reveal potential therapeutic targets. The development of a cellular model efficiently recapitulating necroptosis, while conveniently enabling different types of somatic cell genetic screens, offered the opportunity of attempting a comprehensive survey of human genes involved in the process, both in terms of positive and negative regulation.

Interestingly, we found that deletion of *FADD* rendered KBM7 cells sensitive to necroptotic cell death upon treatment with SMAC mimetic as single agent. In addition to sensitizing cells to the cytotoxic activity of TNFα, SMAC mimetic treatment is known to trigger TNFα production in certain cell lines(51–54). In line with the notion of autocrine TNFα secretion, phosphorylation of MLKL in KBM7 *FADD*^-^ cells treated with SMAC mimetic alone occurred in a delayed manner as compared to TNFα-treated samples (Supplementary Figure 1g). SMAC mimetics are considered promising anti-cancer agents and birinapant, having shown efficacy as a single agent against some tumors and leukemias(55, 56), is currently being evaluated in clinical trials for several malignancies. Interestingly, we found that targeting of *TNFRSF1B* (encoding for TNFR2) rendered cells more resistant towards SMAC mimetic-mediated cell killing. In contrast to TNFR1, TNFR2 can be fully activated only by recognition of membrane-bound, but not soluble TNFα(57, 58). Activation of TNFR2 has been shown to sensitize macrophages for TNFR1-mediated necroptosis through TNFR2-induced depletion of TRAF2-cIAP1/2 complexes and autocrine production of TNFα(59). The fact that TNFR2 was identified here in a pooled screen setting points to a cell-intrinsic function of TNFR2 in enhancing cell death. Intriguingly, a recent study found the birinapant to enhance cell killing via the membrane bound form of TNFα(60). In light of our findings, it will thus be interesting to evaluate a possible role for TNFR2 in these settings and whether sensitivity to SMAC mimetic treatment is correlated with TNFR2 expression. Among the other genes conferring resistance under the strong necroptosis-inducing conditions used, it is tempting to speculate that some of the transcription factors identified are directly or indirectly involved in transcriptional regulation of the different pathway components or TNFα itself. Indeed, the basal transcription factor SP1 has been shown to be recruited to the TNF promoter(61, 62) and to mediate RIPK3 expression(63).

The prominent identification across multiple screens of the *SLC39A7*, to date not linked to TNFα signaling, was of particular interest, considering the instrumental role of SLC transporters in influencing essential physiological processes by regulating metabolic fluxes between the environment and intracellular compartments(64–66). *SLC39A7* controls zinc efflux from the ER to the cytoplasm and its absence has been linked to disturbed zinc homeostasis and ER stress(29–31, 36). We here experimentally link the resistance to ligand-triggered cell death induction observed in *SLC39A7*^-^ cells to a trafficking defect of the respective receptors. This is reminiscent of the Notch trafficking abnormalities described for mutants in *Catsup*, the *Drosophila* ortholog of *SLC39A7*(36). Whereas we found that basic glycosylation of TNFR1 occurs in *SLC39A7*^-^ cells, the receptor does not transit to the Golgi apparatus, indicating that proper folding is not attained. Several of the folding factors in the ER, including protein disulfide isomerases (PDIs) and the chaperones calnexin and calreticulin, rely on zinc as a cofactor and could therefore be affected by SLC39A7 deficiency. Indeed, PDI activity can be inhibited by high zinc concentration(31). Importantly, reconstitution of *SLC39A7*^-^ cells with *wildtype* SLC39A7, but not with mutants predicted to impair its transport activity(40), restored all affected cellular functions indicating that, firstly, the perturbations induced by SLC39A7 deficiency are reversible and, secondly, that substrate-transport is required for its function. Remarkably, the trafficking of specific membrane receptors is differentially affected by SLC39A7 deficiency. We did not identify obvious differences in their posttranslational processing with regard to glycosylation and disulfide bridge formation, and thus the definition of the molecular basis for this divergence remains an intriguing question to be addressed in future studies.

Our data show that SLC39A7 is the prominent SLC affecting TNFR1 and Fas receptor signaling and suggest a largely non-redundant role in ER homeostasis. These are central processes involved in patho-physiology of both the innate and adaptive immune compartment and beyond. It is tempting to speculate that loss-of-function or hypomorphic mutations in SLC39A7 may result in genetic diseases associated with perturbed zinc and ER homeostasis as well as TNFR, FAS and possibly other immune receptor signaling. Indeed, defective FASL-induced lymphocyte apoptosis underlies a group of primary immunodeficiencies (PIDs) denoted as autoimmune lymphoproliferative syndrome (ALPS) and our screen with FASL retrieved SLC39A7 amongst genes previously identified in ALPS- and ALPS-like PIDs such as FAS, FADD, and CASP8 (Supplementary Figures 4b)(67). Similarly, perturbation of the ER compartment can also result in PIDs, as highlighted by the primary antibody deficiency due to plasma cell defects observed in patients with mutations in the Sec61 translocon subunit SEC61A1(68). Conversely, mutations in *TNFRSF1A,* encoding TNFR1 can lead to the autoinflammatory disorder TNFR-associated periodic syndrome (TRAPS)(69).

Complementing the loss-of-function approaches, we used CRISPR-based gene activation technology to perform genome-scale overexpression screens in analogous settings. In order to identify genes conferring a strong protective effect, we selected conditions ensuring a high selective pressure and applied stringent criteria for hit selection, i.e. the significant enrichment of at least two independent sgRNAs. Our data highlights a number of candidate genes conferring resistance to necroptotic cell death and revealed a novel role for TNIP1 in inhibiting necroptosis. TNIP1 has been shown to negatively regulate multiple inflammatory pathways through, in particular, its NF-kB inhibitory and anti-apoptotic activities(42, 43, 45). We demonstrated that TNIP1 overexpression prevented TNFα-induced phosphorylation of RIPK1 and of the downstream effector MLKL via a ubiquitin-binding dependent mechanism, while NF-kB and MAPK signaling was largely unaffected. This expands the role of TNIP1 in controlling TNFα-induced cell death pathways beyond the proposed role in inhibiting apoptosis by interfering with the recruitment of caspase-8 by FADD(45). Considering the association of TNIP1 with multiple inflammatory and autoimmune diseases(42) as well as the TNIP1-dependent phenotypes observed in animal models(46–49, 70), it will be of importance to investigate the relative contribution of TNIP1-dependent necroptosis inhibition in these contexts. Therefore, our work, identifying a novel inhibitory function of TNIP1 on necroptosis, might contribute to a better understanding of these molecular mechanisms and their associated pathologies. In line with our results, a recent study, published while this manuscript was in preparation, showed that TNIP1-deficiency sensitizes cells to TNFα-induced necroptosis by affecting RIPK1 activation(71). Upon TNFα treatment, TNIP1 is recruited to the TNF receptor signaling complex by the action of the Met1-ubiquitylating complex LUBAC, resulting in the engagement of the deubiquitinating enzyme A20, which targets RIPK1 and limits its activation(71). In summary, we presented here complementary genome-scale loss- and gain-of-function screens in a novel, genetically engineered cellular model for necroptosis, and mechanistically validated the role of SLC39A7 and TNIP1 in death receptor trafficking and signaling. Highlighting multiple novel candidate regulators affecting TNFα responses and necroptotic cell death, our work provides the basis for future studies aiming at defining the individual contributions in these central immune processes and exploring the role in associated pathological conditions.

## Acknowledgments

We thank all members of the Superti-Furga laboratory for discussions and feedback, Georg E. Winter for support with the haploid genetic screens, Hawwa O. Oshafu for technical help, the Proteomics Facility for the proteomics analyses and the Biomedical Sequencing facility for the NGS sequencing. This work was supported by the Austrian Academy of Sciences, the European Research Council (ERC) under the European Union’s Horizon 2020 research and innovation program (grant agreement n° 695214), EMBO long-term and Marie Sklodowska-Curie fellowships to M.R. (ALTF 1346-2011, IEF 301663), Marie Sklodowska-Curie fellowship to E.G. (MSCA-IF-2014-661491), Austrian Science Fund (FWF) Lise Meitner Program Fellowship to K.L.W. (FWF M-1809), and Austrian Science Fund grant for J.W.B. (FWF SFB F47). Plasmids obtained through Addgene were a gift from Feng Zhang, David Root and Didier Trono.

## Author contributions

A.F., M.R., K.L.W., K.B. and G.S-F. designed research; A.F., M.R., K.L.W., E.G., S.S., M.B. and J.K. performed research; E.G. generated reagents and provided scientific insight.; A.C-R. and F.S. analyzed genetic screening data; J.W.B., L.X.H. and K.H. provided reagents and gave experimental advice; A.F., M.R., K.L.W. and G.S-F. analyzed and interpreted the data; A.F., M.R. and G.S-F. wrote the paper.

## Competing financial interests

The authors declare no competing financial interest.

## MATERIALS AND METHODS

### Cell lines and reagents

HEK293T were obtained from ATCC (Manassas, VA, USA). KBM-7 and HAP1 were obtained from T. Brummelkamp. HeLa were provided by M. Hentze, Jurkat E6.1 by W. Ellmeier. KBM7 *TNFR1*^-^ cells were obtained from Haplogen (P00371D06). Cells were cultured in DMEM (Gibco, Grand Island, NY, USA), RPMI (Gibco) or IMDM medium (Gibco) supplemented with 10% (v/v) FBS (Gibco) and antibiotics (100 U/mL penicillin and 100 mg/mL streptomycin) (Gibco). Cell lines were checked for mycoplasma by PCR or ELISA. The reagents used were as follows: recombinant human TNF-α (300-01A, Peprotech), SMAC mimetic birinapant (S7015, Selleck Chemicals), FasLigand (ALX-522-020-C005, Enzo), recombinant human TRAIL (310-04, Peprotech), z-VAD-FMK (AG-CP3-0002, Adipogen), necrosulfonamide (480073, Merck Millipore), necrostatin-1 (N9037, Sigma-Aldrich), RIPK1 Inhibitor II 7-Cl-ONec-1 (Nec-1s) (504297, Merck Millipore), PNGase F (P0704, NEB), Endo H (P0702, NEB), MG-132 (S2619, Selleckchem), Thapsigargin (P9033, Sigma-Aldrich), Brefeldin A (1231, Tocris), Tunicamycin (T7765, Sigma Aldrich), and doxycycline (D9891, Sigma-Aldrich, St.Louis, MO, USA).

### Antibodies

Antibodies used were phospho-IkBa (Ser32/36) (9246, Cell Signaling), phospho-SAPK/JNK (Thr183/Tyr185) (9251, Cell Signaling), SAPK/JNK (9252, Cell Signaling), IkBa (SC-371, Santa Cruz), phospho-IKKα/β(Ser176/180) (2697, Cell Signaling), Phospho-p38 MAPK (Thr180/Tyr182) (9215, Cell Signaling), Phospho-NF-κB p65 (Ser536) (3033, Cell Signaling), tubulin (ab7291, Abcam), actin (AAN01-A, Cytoskeleton), BiP (610978, BD Biosciences), CHOP (MA1-250, ThermoFisher), SLC39A7 (19429-1-AP, Proteintech), TNFR1 (sc-8436, Santa Cruz), RIPK1 (610458, BD Bioscience), RIPK3 (12107, Cell Signaling), GFP (sc-69779, Santa Cruz), FADD (610399, BD Biosciences), phospho-MLKL (Ser385) (ab187091, Abcam), phospho-RIPK1 (S166) (65746, Cell Signaling), phospho-RIPK3 (S227) (ab209384, abcam), cleaved Caspase-3 (Asp175) (9661, Cell Signaling), V5 (R960-25, Invitrogen or ab9116, abcam), PDIA2 (ab2792, abcam), TNIP1/ABIN-1 (4664, Cell Signaling), cIAP2/BIRC3 (3130, Cell Signaling) and LAMP1 (ab25630, abcam). The secondary antibodies used were goat anti-mouse HRP (115-035-003, Jackson ImmunoResearch), goat anti-rabbit HRP (111-035-003, Jackson ImmunoResearch), Alexa Fluor 680 goat anti-mouse (A-21057, Molecular probes) and IRDye 800 donkey anti-rabbit (611-732-127, Rockland).

### Plasmids and cloning

CRISPR/Cas9-based knockout cell line generation was performed using pX330-U6-Chimeric_BB-CBh-hSpCas9 (Addgene plasmid #42230), pLentiCRISPRv2 (Addgene plasmid #52961), or pLentiCas9-BlastR (Addgene plasmid #52962) and pLentiGuide-PuroR (Addgene plasmid #52963). To enable color tracing of targeted cells in competition assays, an IRES-GFP or IRES-mCherry fragment was inserted into pLentiGuide-PuroR creating LGPIG (pLentiGuide-PuroR-IRES-GFP) and LGPIC (pLentiGuide-PuroR-IRES-mCherry) using standard cloning techniques. CRISPR cloning was performed as described elsewhere(23, 72). In brief, sgRNAs for KO generation were designed using crispr.mit.edu or CHOPCHOP(73). Oligonucleotides containing *Bsm*BI restriction site-compatible overhangs were annealed, phosphorylated and ligated into pX330-U6-Chimeric_BB-CBh-hSpCas9, pLentiCRISPRv2, LGPIG or LGPIC using standard cloning techniques and sequence verified using Sanger sequencing. sgRNA targeting *Renilla luciferase* coding sequence (sg*Ren*) was used as a negative control. The sequences of sgRNAs used in this study are listed in Supplementary Table 3. sgRNAs are labeled by target gene name followed by the sequence position of the guide in the corresponding mRNA sequence. Sequences of sgRNAs for SAM-mediated transcriptional activation of TNIP1 and BIRC3 were extracted from hits of the genome-scale library screen and cloned into pLenti-sgRNA(MS2)-zeo backbone (Addgene plasmid #61427). sgRNA targeting *Renilla luciferase* coding sequence (sg*Ren*) was used as a negative control and cloned into pLenti-sgRNA(MS2)-zeo-IRES-GFP backbone. SLC39A7 (Clone ID: 3345970, Dharmacon, GE Healthcare) and TNIP1 (HsCD00042052, Harvard Medical School plasmID repository) coding sequence was amplified and subcloned into vector pDONR221 using Gateway technology (Invitrogen, Grand Island, NY, USA). Point mutations were introduced by site-directed mutagenesis (InvivoGen). Following sequence verification, cDNAs were transferred into Gateway-compatible expression vectors pLIX403 (Addgene plasmid #41395) for confocal microscopy experiments, pLX304 (Addgene plasmid #25890) for SLC39A7 cDNA rescue experiments or pLX302 (Addgene plasmid #25896) for TNIP1 cDNA overexpression. Retroviral packaging plasmids pGAG-POL and pVSV-G were obtained from T. Brummelkamp, and pADVANTAGE from Promega (E1711). Lentiviral packaging plasmids psPAX2 (plasmid #12260) and pMD2.G (plasmid #12259) were from Addgene.

### Cell line generation

HEK293T cells were transiently transfected with pGAG-POL, pVSV-G, pADVANTAGE and retroviral expression vectors, or psPAX2, pMD2.G and lentiviral expression vectors using Polyfect (301105, QIAGEN). After 24h the medium was replaced with fresh medium. The virus-containing supernatant was harvested 48h later, filtered (0.45 μm), supplemented with 8 μg/ml protamine sulfate (P3369, Sigma-Aldrich) and added to 40-60% confluent target cells. Suspension cell lines were subjected to spinfection (2000 rpm, 45 min, RT). 24h after infection, the medium was replaced with fresh medium supplemented with the respective antibiotics to select for infected cells. Where indicated, target gene expression was induced by adding 2 µg/ml doxycycline. For generation of KBM7 *FADD*^-^ cells pX330-U6-Chimeric_BB-CBh-hSpCas9 harboring a *FADD*-targeting sgRNA was electroporated into KBM7 cells using Nucleofector^TM^ Technology (Lonza). In brief, 2×10^6^ cells were spun down, resuspended in 100 µl Nucleofection solution (Amaxa Cell Line Nucleofector Kit V, Lonza), and transferred to an electrode cuvette. After addition of 1µg of target vector DNA and electroporation (program Y-005), cells were immediately resuspended in 700 µl medium without antibiotics and transferred to a culture vessel with pre-warmed medium. Single cell knockout clones were derived by limiting dilution from KBM7 *FADD-Cas9* cells transduced with LGPIG sg*SLC39A7*_618 or KBM7 *FADD-*transduced with pLentiCRISPRv2 sg*SLC39A7_*618.

### Haploid genetic screens and deep sequencing analysis

Haploid genetic screening was conducted as described previously(19, 74). In brief, gene-trap virus was produced by transient transfection of low passage, subconfluent HEK293T cells with the gene-trap plasmid and packaging plasmids pGAG-POL, pVSV-G and pADVANTAGE using Lipofectamine 2000 (Thermo Fisher Scientific). Every 24h for three consecutive times the virus-containing supernatant was collected and replaced with fresh medium. Virus was concentrated via ultracentrifugation and used to mutagenize 1×10^8^ KBM7 cells by spinfection. Mutagenized cells were then expanded and directly used for genetic screens, while 1×10^8^ gene-trapped cells were harvested as unselected control population. For screens, a total number of 1×10^8^ gene-trapped cells/screen was selected with the indicated stimuli (TRAIL 100 ng/ml, FASL 250 ng/ml) in 96-well plates (1×10^5^ cells/well in 100 µl). After 12 days, resistant clones were pooled, cleaned up using Lonza lymphocyte separation buffer (17-829E), expanded to a total number of 3-6×10^7^ cells, and genomic DNA was isolated.

To map insertions by next generation sequencing, retroviral insertion sites in necroptosis screen samples and unselected reference pools were recovered via linear amplification-mediated (LAM)-PCR, and FASL and TRAIL screen samples were digested with NlaIII and MseI and circularized, followed by inverse PCR. Samples were loaded onto an Illumina HiSeq 2000 machine using custom sequencing primer (18) and sequenced 50 base pair single-end. The obtained sequences in the FASTQ data file were mapped to the human reference genome hg19 (UCSC hg19 build) using bowtie2 (version 2.2.4) with default parameters. Sequencing reads with multiple alignment as well as reads with low mapping quality (MAPQ<20) were discarded. Duplicate reads were marked and discarded with Picard (version 1.118). Insertions 1 or 2 base pairs away from each other were removed to avoid inclusion of insertions due to mapping errors. Each uniquely mapped read corresponds to an insertion site and was annotated with gene build GRCh37.p13 (ENSEMBL 75 - release February 2014) using bedtools (version 2.10.1) and custom scripts. Only insertions that are predicted to be disruptive to gene function were further tested for significant enrichment. The significance of enrichment of insertions in a given gene was calculated by comparing the number of insertions in the selected populations with the unselected control data set by applying a one-sided Fisher’s exact test. P-values were adjusted for false discovery rate (FDR) using Benjamini-Hochberg procedure (5% threshold was considered significantly enriched). Screen results were visualized using Circos (version 0.66) software (75), summary bubble plots and gene-trap insertion plots were generated via custom scripts using R statistical environment.

### CRISPR/Cas9-based genetic screens

The SLC KO CRISPR/*Cas9* library used is described in detail in Girardi et al. (in preparation). Briefly, a CRISPR/*Cas9* library targeting 388 SLC genes with six sgRNAs per gene, together with a set of 120 sgRNAs targeting 20 genes essential in KBM7 and HAP1 cells(35), and a set of 120 non-targeting sgRNAs was cloned by Gibson cloning in the pLentiCRISPRv2 lentiviral vector. Viral particles were generated by transient transfection of low passage, subconfluent HEK293T cells with the SLC-targeting library and packaging plasmids pGAG-POL and pVSV-G using PolyFect (Qiagen). After 24h the media was changed to fresh IMDM media supplemented with 10% FCS and antibiotics. The viral supernatant was collected after 48h, filtered and stored at -80°C until further use. The supernatant dilution necessary to infect KBM7 *FADD*^-^ cells at a MOI (multiplicity of infection) of 0.2-0.3 was determined by puromycin survival after transduction as described in(76). KBM7 *FADD*^-^ cells were infected in duplicates with the SLC KO library at high coverage (1000x) and after selection for 7 days with puromycin (0.5 μg/ml) an initial sample was collected to control for library composition. Cells were then treated with a set of stimuli (KBM7 *FADD^-^*: TNFα 50ng/ml, 12 days; TNFα 10ng/ml, 12 days; SMAC mimetic 5 μM, 12 days; SMAC mimetic 1 μM, 12 days; KBM7 *wildtype*: TRAIL 100 ng/ml, 14 days; FASL 100 ng/ml, 10 days; HAP1 *wildtype*: TRAIL 100 ng/ml, 9 days; FASL 100 ng/ml, 7 days) and cell samples collected from both treated and control samples. Genomic DNA was extracted using the DNAeasy kit (QIAGEN) and the cassette containing the sgRNA sequence amplified with two rounds of PCRs following the procedure described in(76).

The amplified samples were sequenced on a HiSeq3000/4000 (Illumina), followed by processing with a custom analysis pipeline (see next section).

Genome-scale SAM screen was performed following an analogous procedure as previously described (23). In brief, KBM7 *FADD*^-^ cells were transduced first with lenti MS2-P65-HSF1 (Addgene plasmid #61426; selection Hygromycin 800 μg/ml) and subsequently with lenti dCAS-VP64 (Addgene plasmid #61425, selection Blasticydin 20 μg/ml) to generate KBM7 *FADD*^-^ SAM cells. The Human CRISPR/Cas9 Synergistic Activation Mediator (SAM) pooled sgRNA library (Addgene #1000000057) was amplified and packaged into lentivirus following the same procedure described above for the SLC KO CRISPR/Cas9 library. After viral titration, 2x10^8^ KBM7 *FADD*^-^ CRISPR-SAM cells were infected at 0.3 MOI and selected 48h later with zeomycin (150 μg/ml) for 8 days. 5x10^6^ cells were either harvested (time 0 control), treated in duplicate with two doses of TNFα (10 or 100 ng/ml) or SMAC mimetic (0,1 or 1 μM) or left untreated (untreated control). After 72h cells were washed, expanded for 7 days before isolation of live cells by Lymphoprep (Stemcell technologies). Genomic DNA was extracted using the DNAeasy Blood & Tissue kit (QIAGEN) and the cassette containing the sgRNA sequence amplified using NEBnext High Fidelity 2X Master Mix (New England Biolabs) in 36 single-step PCR reactions of 23 cycles per sample with primers introducing sequencing adaptors and barcodes (described in (23)). PCR products were pooled, purified and gel extracted. The amplified samples were then quantified, pooled (maximum of 8 samples per sequencing lane) and sequenced on a HiSeq3000/4000 (Illumina), followed by processing with a custom analysis pipeline (see next section).

### Analysis of CRISPR screens

Sequences of sgRNAs were extracted from RNA-Seq reads, matched against the original sgRNA library index and counted using an in-house Python script. To compensate for the noise and off target action of sgRNAs inherent to CRISPR screening approaches, we used a two-step differential abundance analysis. In a first step, differential abundance of sgRNAs was estimated with DESeq2(77) using a two-factor design that accounts for both time and treatment variables. Subsequently, significantly enriched sgRNAs (adjusted p-value ≤ 0.05) were sorted by log2FoldChange and aggregated to genes using Gene Set Enrichment Analysis(78), only considering genes with at least two significantly enriched sgRNAs.

### T7 Endonuclease assay

To determine *FADD* targeting efficiency, 1-3x10^6^ cells were spun down, washed in 1x PBS and genomic DNA isolated using the DNeasy Blood & Tissue Kit (QIAGEN) according to the manufacturer’s instructions. The CRISPR/*Cas9* targeting site-containing locus was PCR-amplified (fw primer: GAGCTGACCGAGCTCAAGTTCCTAT; rv primer: CAAATCAAACCCGGCAAAGG; product size: 342 bp). The PCR products were denatured at 95°C for 2 min and re-annealed by ramping from 95-85°C at -2°C/s and from 85-25°C at -0.1°C/second. The annealed PCR products were digested with T7 Endonuclease I (NEB, M0302) for 20 min at 37°C, control digests without T7 Endonuclease I served as negative control. The fragmented PCR products were analysed by agarose gel electrophoresis.

### Immunoblotting

Whole cell extracts were prepared using Nonidet-40 lysis buffer (50 mM HEPES pH 7.4, 250 mM NaCl, 5 mM EDTA, 1% NP-40, 10 mM NaF and 1 mM Na3VO4 or Halt phosphatase inhibitor cocktail (ThermoScientific), one tablet of EDTA-free protease inhibitor (Roche) per 50 ml) for 10 min on ice. Lysates were cleared by centrifugation (13000 *rpm*, 10 min, 4°C). The proteins were quantified with BCA (Pierce, Grand Island, NY, USA) or Bradford assay (Bio-Rad, Hercules, CA, USA). Cell lysates were resolved by SDS-PAGE and transferred to nitrocellulose membranes Protran BA 85 (GE Healthcare, Little Chalfont, UK). The membranes were immunoblotted with the indicated antibodies. Bound antibodies were visualized with horseradish peroxidase–conjugated secondary antibodies using the ECL Western blotting system (Thermo Scientific, Waltham, MA, USA) or Odyssey Infrared Imager (LI-COR, Lincoln, NE, USA).

### Cell viability assays

Cells were seeded in 96-well plates at the appropriate cell density. For cell death induction, cells were incubated with the indicated compounds as stated or overnight (14-16h). Cell viability was determined using CellTiter Glo Luminescent Cell Viability Assay (Promega, Fitchburg, WI, USA) according to the instructions provided by the manufacturer. Luminescence was recorded with a SpectraMax M5Multimode plate reader (Molecular Devices, Sunnyvale, CA, USA). Data were normalized to values of untreated controls.

### Flow cytometry

For flow cytometric analyses, cells were blocked with PBS, 10% FBS for 10 min on ice and subsequently stained with PE-labeled mouse anti-human TNFR1 (R&D Systems, FAB225P), PE-Cy^TM^7-labeled mouse anti-human CD95 (BD Biosciences, 561633), PE-labeled mouse anti-human TRAIL-R1/DR4 (eBioscience, 12-6644-41), PE-labeled mouse anti-human TRAIL-R2/DR5 (eBioscience, 12-9908-41), APC-labeled mouse anti-human C3AR (BioLegend, 345805), PE-labeled mouse anti-human C5L2 (BioLegend, 342403), FITC-labeled mouse anti-human CD11a (eBioscience, 11-0119-41), PE-labeled mouse anti-human CD123 (eBioscience, 12-1239-42), PE-Cyanine5-labeled mouse anti-human CD14 (eBioscience, 15-0149-41), APC-labeled mouse anti-human CD31 (eBioscience, 17-0319-42), Cy^TM^7-labeled mouse anti-human CD34 (BD Pharmingen, 560710), PerCP-Cyanine5.5-labeled mouse anti-human CD4 (eBioscience, 45-0049120), PerCP-Cyanine5.5-labeled mouse anti-human CD44 (BD Pharmingen, 560531), PerCP-Cyanine5.5-labeled mouse anti-human CD45 (eBioscience,45-0459-42), PE-labeled mouse anti-human CD54 (BD Pharmingen, 560971), PE-labeled mouse anti-human CD55 (BD Pharmingen, 561901), or PerCP-Cyanine5.5-labeled mouse anti-human HLA-ABC (BD Pharmingen, 555554). All stains were performed in FACS buffer (PBS, 10% FBS) for 30 min in the dark at either RT or 4°C, followed by two washing steps. Samples were analyzed on an LSR Fortessa (BD Biosciences) or FACSCalibur (BD Biosciences). Dead cells were excluded by forward- and side-scatter, and data analysis was performed using FlowJo software version 7.6.3 (Tree Star Inc., Ashland, OR, USA).

### Flow cytometry-based multi-color competition assay (MCA)

To analyze the long-term cellular response upon treatment with different cell death inducers in CRISPR/*Cas9*-based knockout cell competition experiments, cell populations were marked with a fluorescent reporter coupled to individual sgRNAs, with sgRNA targeting *Renilla luciferase* (sg*Ren*) serving as control. sg*Ren*-mCherry^+^ reporter cells (LGPIC lentiviral sgRNA vector) were mixed with sg*Ren*-GFP^+^ control or target gene sgRNA-GFP^+^ cells (LGPIG lentiviral vector) in a 1:1 ratio. The mixed cell populations were treated with the cell death inducers indicated or left untreated as reference. After 14 days, the respective percentage of mCherry^+^ and GFP^+^ was determined by flow cytometry, gating on viable cells (FSC/SSC).

### Confocal microscopy

HeLa cells inducibly expressing V5-tagged SLC39A7 were plated on glass coverslips and expression induced by addition of 2 µg/ml doxycycline. After 24h, cells were washed with 1x PBS, fixed (PBS, 4% formaldehyde) and permeabilized (PBS, 0.3% saposin, 10% FBS). Subsequently, coverslips were incubated with anti-V5 (Invitrogen, R960-25) and anti-Calreticulin (abcam, ab2907) for 1h at RT in PBS, 0.3% saposin, 10% FBS. After three washes, coverslips were stained with goat anti-mouse AlexaFluor 568 (Invitrogen, A-11004) and anti-rabbit AlexaFluor488 (Invitrogen, A-11008) antibodies followed by DAPI staining. Coverslips were washed 3 times and mounted on glass slides using ProLong Gold (Invitrogen, P36934). Images were taken with a Zeiss Laser Scanning Microscope (LSM) 700. Images were taken with a 63× oil immersion objective (na 1.4) and exported from lsm to tiff files.

### Triton X-114 phase separation for enrichment of membrane proteins

Triton X-114 phase separation was performed as described previously (79). Briefly, 60-75×106 cells were harvested and washed once with PBS. Cells were then resuspended in 500 μL PBS and 100 μL 6 % pre-condensed Triton X-114, mixed by inversion and incubated for 15 min on ice. After centrifugation for 1 min at 13000 rpm, the supernatants were transferred to new tubes and incubated for 5 min at 37°C to induce phase separation. The upper aqueous phase was transferred to a new tube. To wash, 500 μL PBS were added to the detergent phase and 100 μL 6 % Triton X-114 to the aqueous phase, followed by incubation for 5 min on ice and 5 min at 37°C. Samples were centrifuged as before and the initial phases were kept for further processing. Proteins in the detergent phase were precipitated by adding 450 µl PBS, 500 μL methanol and 125 μL chloroform followed by vortexing. After centrifugation for 4 min at 13000 rpm, 750 µl of the supernatant were removed, 400 µl methanol added and mixed by pipetting. Samples were centrifuged for 1 min at 13000 rpm, the supernatant removed and the pellets dried at RT. Subsequently, the proteins were resuspended by sonication in 100 µl SDS-PAGE sample buffer without glycerol and bromophenol blue. Samples were then either subjected to immunoblot analysis or further prepared for mass spectrometric analysis via the FASP (filter aided sample preparation) method as described previously(80).

### Reversed-phase liquid chromatography mass spectrometry (LCMS)

Mass spectrometry was performed on a hybrid linear trap quadrupole (LTQ) Orbitrap Velos mass spectrometer (ThermoFisher Scientific, Waltham, MA) using the Xcalibur version 2.1.0 coupled to an Agilent 1200 HPLC nanoflow system (dual pump system with one pre-column and one analytical column) (Agilent Biotechnologies, Palo Alto, CA) via a nanoelectrospray ion source using liquid junction (Proxeon, Odense, Denmark). Solvents for LCMS separation of the digested samples were as follows: solvent A consisted of 0.4% formic acid in water and solvent B consisted of 0.4% formic acid in 70% methanol and 20% isopropanol. From a thermostatic microautosampler, 8 μL of the tryptic peptide mixture were automatically loaded onto a trap column (Zorbax 300SB-C18 5 μm, 5 × 0.3 mm, Agilent Biotechnologies) with a binary pump at a flow rate of 45 μL/min. 0.1% TFA was used for loading and washing the precolumn. After washing, the peptides were eluted by back-flushing onto a 16 cm fused silica analytical column with an inner diameter of 50 μm packed with C18 reversed phase material (ReproSil-Pur 120 C18-AQ, 3 μm, Dr. Maisch, Ammerbuch-Entringen, Germany). The peptides were eluted from the analytical column with a 27 min gradient ranging from 3 to 30% solvent B, followed by a 25 min gradient from 30 to 70% solvent B and, finally, a 7 min gradient from 70 to 100% solvent B at a constant flow rate of 100 nL/min. The analyses were performed in a data-dependent acquisition mode using a top 10 collision-induced dissociation (CID) method. Dynamic exclusion for selected ions was 60s. A single lock mass at m/z 445.120024 was employed. The maximal ion accumulation time for MS in the orbitrap and MS2 in the linear trap was 500 and 50 ms, respectively. Automatic gain control (AGC) was used to prevent overfilling of the ion traps. For MS and MS2, AGC was set to 10^6^ and 5,000 ions, respectively. Peptides were detected in MS mode at a resolution of 60,000 (at m/z 400). The threshold for switching from MS to MS2 was 2,000 counts. All samples were analysed as technical, back-to-back replicates.

### Mass spectrometry data processing and analysis

The acquired raw MS data files were processed with msconvert (ProteoWizard Library v2.1.2708) and converted into Mascot generic format (mgf) files. The resultant peak lists were searched against either the human or mouse SwissProt database v2014.07 (40,984 and 24,862 sequences, respectively, including isoforms obtained from varsplic.pl(81) and appended with known contaminants) with the search engines Mascot (v2.3.02, MatrixScience, London, U.K.) and Phenyx (v2.5.14, GeneBio, Geneva, Switzerland)(82). Submission to the search engines was via a Perl script that performs an initial search with relatively broad mass tolerances (Mascot only) on both the precursor and fragment ions (±10 ppm and ±0.6 Da, respectively). High-confidence peptide identifications were used to recalibrate all precursor and fragment ion masses prior to a second search with narrower mass tolerances (± 4 ppm and ±0.3 Da, respectively). One missed tryptic cleavage site was allowed. Carbamidomethyl cysteine and oxidised methionine were set as fixed and variable modifications, respectively. To validate the proteins, Mascot and Phenyx output files were processed by internally-developed parsers. Proteins with ≥2 unique peptides above a score T1 or with a single peptide above a score T2 were selected as unambiguous identifications. Additional peptides for these validated proteins with score >T3 were also accepted. For Mascot and Phenyx, T1, T2, and T3 peptide scores were equal to 14, 28, 10 and 4.2, 4.7, 3.5, respectively (*p*-value <10^-3^). The validated proteins retrieved by the two algorithms were merged, any spectral conflicts discarded and grouped according to shared peptides. A false discovery rate of <1% for protein identifications and <1% for peptides (including the ones exported with lower scores) was determined by applying the same procedure against a database of reversed protein sequences. Commonly-known contaminants including trypsin and keratin were removed. In each condition, four spectral counts were available for the median (two biological replicates and two technical for each). Average spectral counts of the replicates were used to determine log2 fold changes between KBM7 *FADD-SpCas9 SLC39A7*^-^ vs. sg*Ren* control. *p*-values were calculated by a two-sided Welsch Two Sample t-test. Annotations for the GO term “GO:0034976 response to endoplasmic reticulum stress” were obtained from QuickGO (www.ebi.ac.uk/QuickGO). GO term enrichment was computed with R’s topGO package for biological process ontology, using the default algorithm (“weight01”) and the annotation file from geneontology.org as of October 2016. The study set consisted in all proteins identified with a log2 fold change ≥1 and a *p*-value <0.05 (65 proteins). All reviewed human proteins in UniProtKB/Swiss-Prot were used as background population (20,348 proteins). *p*-values were corrected for multiple testing using Benjamini-Hochberg’s procedure (False Discovery Rate). Gene Set Enrichment Analysis was performed using the GSEA software from the Broad Institute with default parameters. The gene list consisted in all proteins identified below a *p*-value of 0.05 (354 proteins), ranked by log2 fold change. The analysis was run against the Hallmarks gene set. Supplementary Table 2 gives results for LC-MSMS analysis, GO term enrichment, and GSEA.

### Experimental design, data plotting and statistical rational

Proteomics experiments were performed as biological replicates (n = 2) and analyzed by LC-MSMS as technical duplicates. CRISPR/*Cas9* library screens were performed in duplicate (n = 2). Flow cytometry-based MCA data are shown as mean value ± s.d. of at least two independent experiments (n ≥ 2) performed in duplicates. Data calculations were performed using Microsoft Excel (Microsoft, Redmond, WA, USA), data plotting and statistical analysis was done using GraphPad Prism 6 (GraphPad Software) if not otherwise stated. A normal distribution of data was assumed and appropriate tests were applied. Visualization of data was performed using R statistical environment.

## SUPPLEMENTARY INFORMATION

**Supplementary Figure 1:**
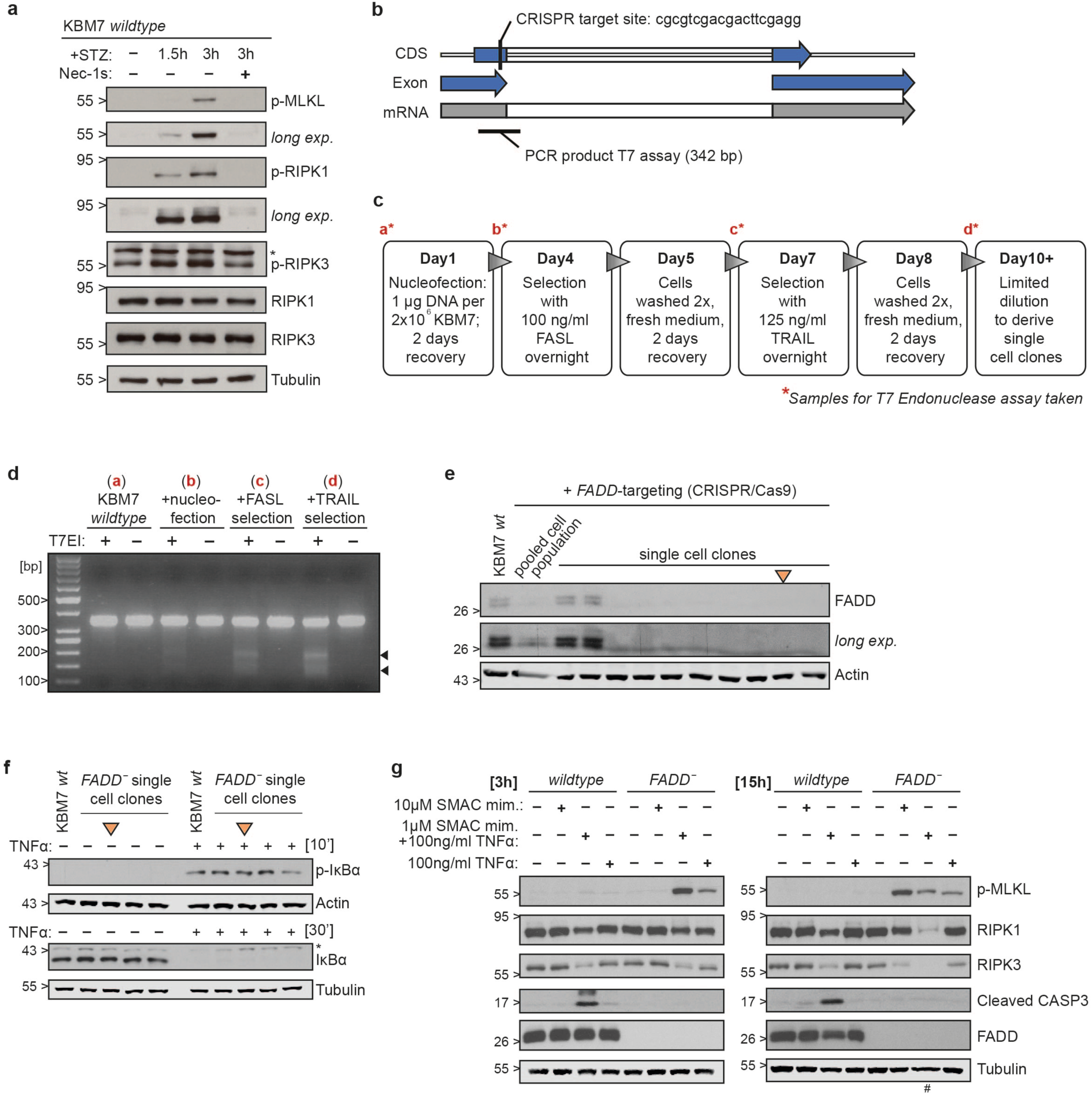
Generation of a *FADD*-deficient KBM7 cell line as novel necroptosis model. (**a**) KBM7 cells were treated with 10 ng/ml TNFα (T), 0.5 µM SMAC mimetic (S) and 20 µM z-VAD-FMK (Z) for the indicated time. Nec-1s (50 µM) treatment serves as control. Cells were lysed subsequently and subjected to immunoblotting with the indicated antibodies, * indicates non-specific band. (**b**) Human genomic *FADD* locus. The CRISPR target site and sequence as well as localization of the PCR product used to monitor editing efficiency by the T7 Endonuclease I assay are indicated. (**c**) Schematic overview for creation and selection of a KBM7 *FADD*^-^ cell line. (**d**) T7 Endonuclease assay to determine *FADD* targeting efficiency. KBM7 cells were harvested at the time points indicated in **c**, and subjected to genomic DNA isolation. The *FADD* locus containing the targeting site was PCR-amplified and the products digested with T7 Endonuclease, followed by agarose gel separation of reaction products. Arrowheads indicate nuclease cleavage products. (**e**) Immunoblot of KBM7 sg*FADD* single cell clones, KBM7 *wildtype* and pooled *FADD*-targeted cell population serve as reference. (**f**) sg*FADD* single cell clones were treated for the time indicated with 10 ng/ml TNFα. Cells were then lysed and subjected to immunoblotting with the indicated antibodies, * indicates non-specific band; KBM7 *FADD*^-^ clone selected for screens is marked in orange in panels **e-f**. (**g**) KBM7 *wildtype* and KBM7 *FADD*^-^ cells were treated for either 3 or 15h with the indicated stimuli. Cells were then lysed and subjected to immunoblotting with the indicated antibodies, ^#^ denotes high number of dead cells/debris in well.

**Supplementary Figure 2:**
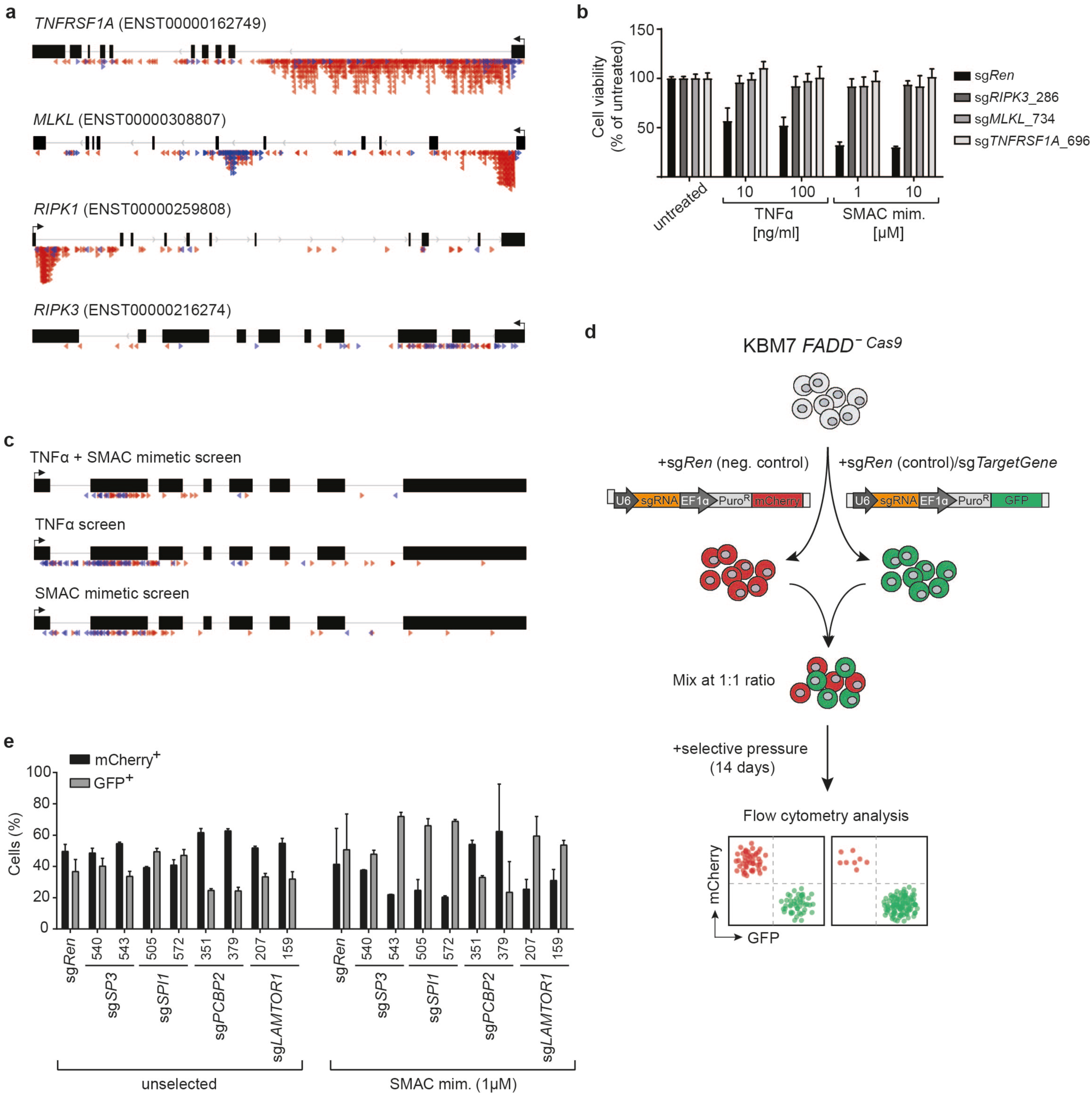
Gene-trap insertion profile and validation of haploid genetic screen hits. (**a**) Genomic location of gene-trap insertions identified in the combined 100 ng/ml TNFα + 1 µM SMAC mimetic screen affecting the known necroptosis mediators *TNFRSF1A*, *MLKL*, *RIPK1*, and *RIPK3*. Sense integrations (red) generate a gene knockout independent of exonic or intronic position. Antisense integration (blue) targeting exons are disruptive, whereas intronic antisense integration has no effect on gene expression in most cases. (**b**) Cell viability in KBM7 *FADD*^-^ lentivirally infected with sgRNAs targeting *RIPK3*, *MLKL*, *TNFR1*, or *Renilla luciferase* (sg*Ren*) as control were treated for 24h with the indicated stimuli. Cell viability was assessed using a luminescence-based readout for ATP (CellTiter Glo). Data represent mean value ± s.d. of two independent experiments performed in triplicates. (**c**) Genomic location of gene-trap insertions (red, sense integration; blue, antisense integration) identified in the indicated haploid genetic screens affecting *SLC39A7*. (**d**) Experimental validation strategy for hits identified in haploid genetic screening approach by CRISPR/*Cas9*-based multicolor competition assay (MCA). KBM7 *FADD^-^ SpCas9* cells were lentivirally infected with sgRNA-expression LentiGuide-PuroR vectors carrying either an mCherry or GFP fluorescent marker, enabling to monitor the respective targeted cell populations using flow cytometry. sg*Ren-*mCherry control cells were mixed with either sg*Ren*-GFP control or gene targeting sgRNA-GFP cells at 1:1 ratio and subjected to selective pressure with the respective indicated stimuli for 14 days. The percentage of GFP^+^ and mCherry^+^ cells in the remaining viable population was then determined by flow cytometry as a measure for survival/growth differences upon selective pressure. (**e**) MCA of KBM7 *FADD^-^ SpCas9* cells transduced with a GFP marker (GFP^+^) and sgRNAs targeting either *SP3, SPI1, PCBP2*, *LAMTOR1*, or *Renilla luciferase* (*sgRen*) as control, against cells transduced with sg*Ren* and an mCherry marker (mCherry^+^). The cell populations were mixed at 1:1 ratio, treated with SMAC mimetic (1 µM), and analyzed after 14 days by flow cytometry. Data represent mean value ± s.d. of two independent experiments performed in duplicates.

**Supplementary Figure 3:**
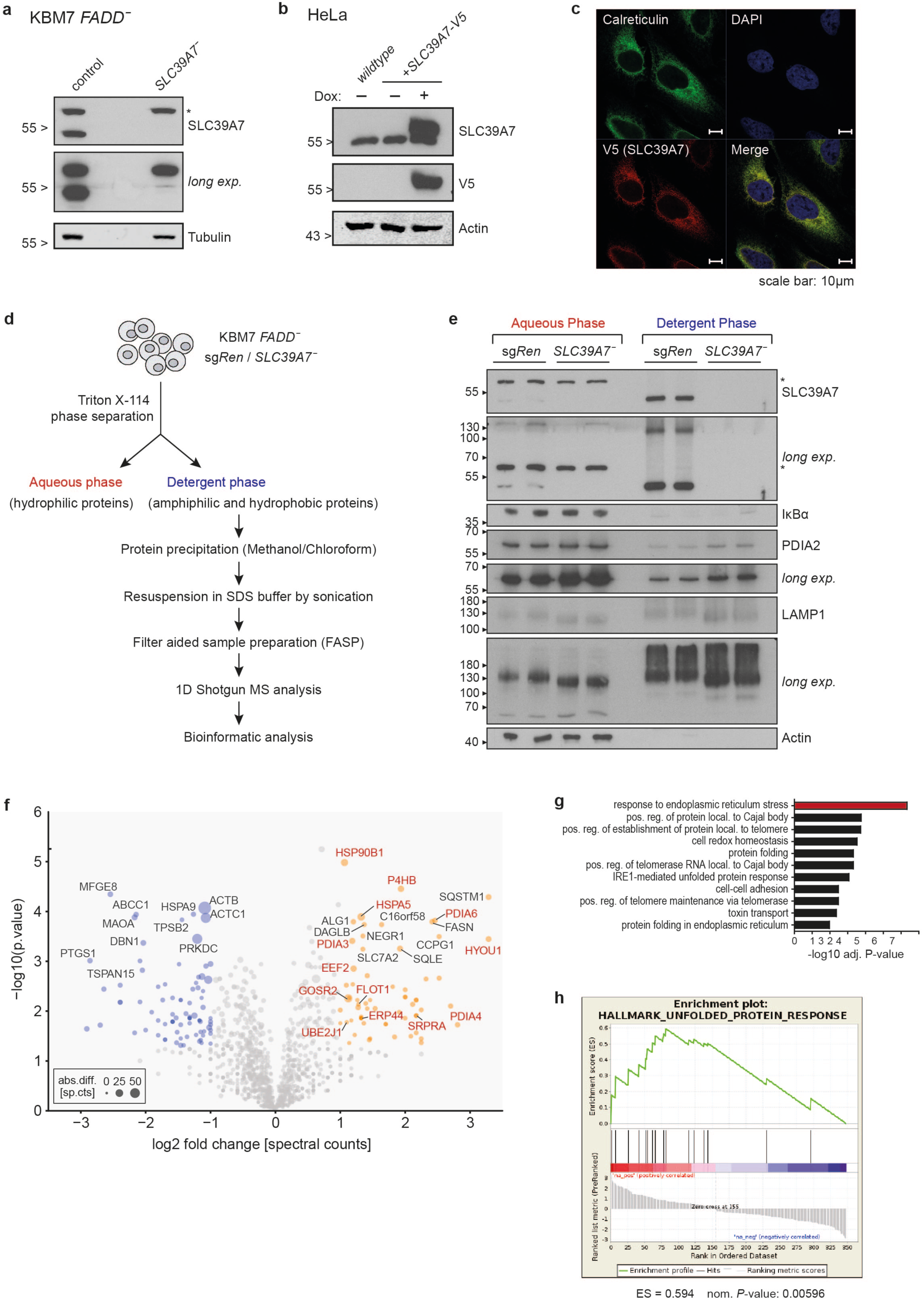
Loss of SLC39A7 leads to an ER stress response. (**a**) Immunoblot of KBM7 *FADD^-^ SpCas9* sg*SLC39A7*_618-derived single cell clone. (**b**) HeLa cells were transduced with an inducible lentiviral vector construct harboring C-terminally V5-tagged *SLC39A7*. Cells were induced with 2 µg/ml doxycycline for 24h and analyzed by immunoblotting using the indicated antibodies. (**c**) Confocal microscopy images of doxycycline-induced HeLa cells expressing SLC39A7-V5 immunostained with anti-V5 and Calreticulin (CALR) antibodies. Nuclei were stained with DAPI. Representative cells are shown; scale bars, 10 µm. (**d**) Workflow for membrane-focused proteomics approach. KBM7 *FADD^-^ SLC39A7*^-^ or sg*Ren* cells were subjected to Triton X-114 phase separation. Proteins recovered in the detergent phase were precipitated using a methanol/chloroform mixture, resuspended in SDS buffer by sonication and prepared for mass spectrometric analysis by filter-aided sample preparation (FASP). (**e**) Immunoblot of Triton X-114 phase-separated KBM7 *FADD^-^ SLC39A7*^-^ or sg*Ren* control samples. Cells were treated as outlined in (**d**) and replicate samples from aqueous and detergent phase immunoblotted using the indicated antibodies, * indicates non-specific band. (**f**) Volcano plot depicting proteins identified by mass spectrometry in KBM7 *FADD^-^ SpCas9 SLC39A7*^-^ cells compared to sg*Ren* control. Log2 fold change in abundance of spectral counts (x-axis) is plotted against enrichment significance (y-axis), bubble size corresponds to absolute difference in spectral counts. Proteins with a *p*-value <10^-3^ are labeled by name; proteins with a log2 fold change in spectral counts >1 belonging to GO term “*response to endoplasmic reticulum stress”* are highlighted in red. Data shown are based on two independent experiments, each analyzed as technical duplicates. (**g**) GO terms enriched (adj. *p*-value <0.01) in upregulated proteins (log2fold change >1) in KBM7 *FADD^-^ SpCas9 SLC39A7*^-^ cells compared to sg*Ren* control. (**h**) Top hit of GSEA using Hallmark gene sets on significantly up- or downregulated proteins (*p*-value <0.05) in KBM7 *FADD^-^ SpCas9 SLC39A7*^-^ cells compared to sg*Ren* control.

**Supplementary Figure 4:**
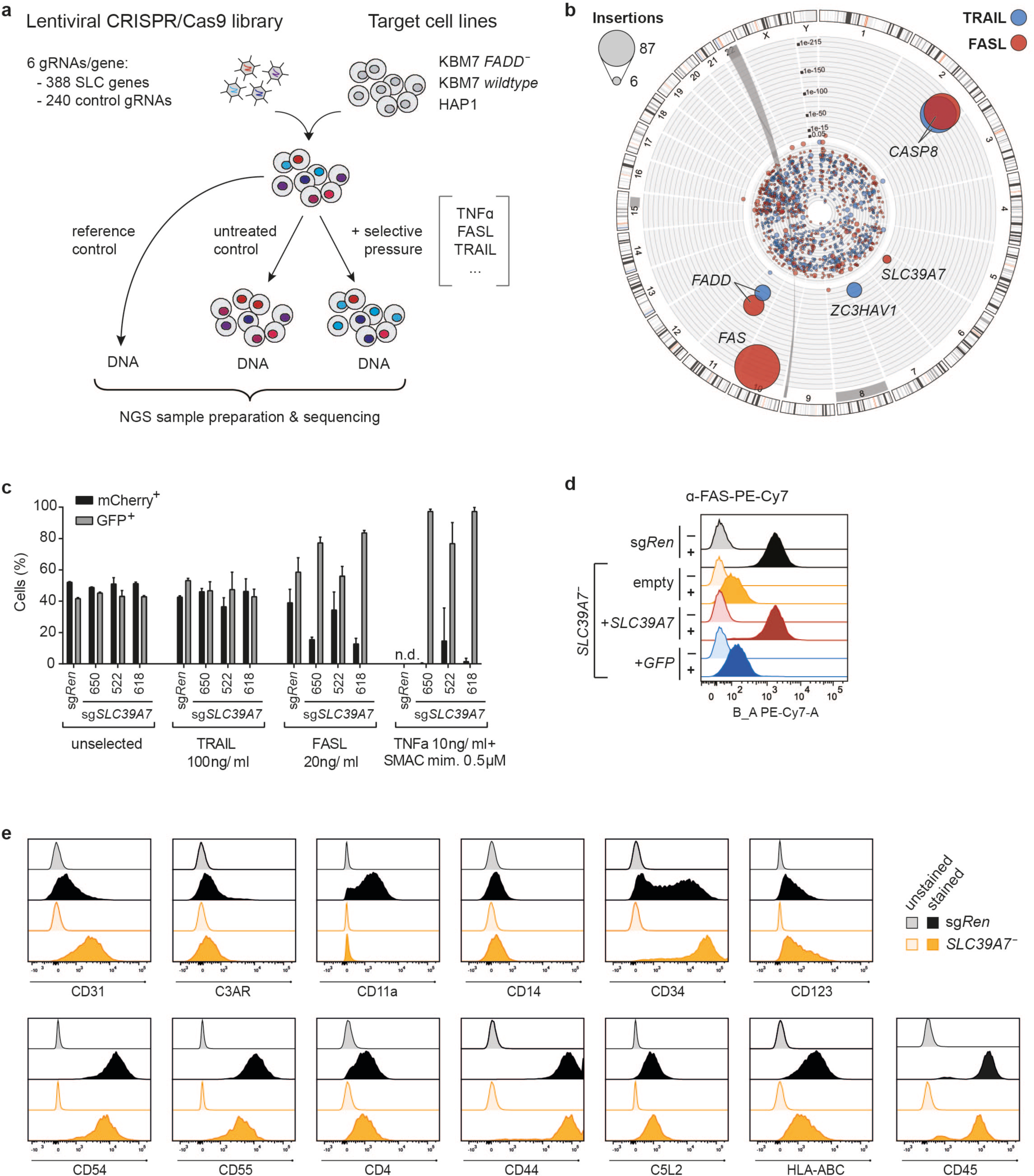
SLC39A7-deficiency differentially affects death receptor trafficking and signaling. (**a**) Schematic overview for CRISPR/*Cas9* library screens. The respective target cell lines were infected with an SLC-specific lentiviral CRISPR/*Cas9* library containing 2345 gRNAs targeting 388 SLC genes (~6 guides/gene) as well as 240 control gRNAs. Library-transduced cell populations were either left untreated or exposed to different cell death inducers. After a short period to allow for recovery and outgrowth, cells were harvested, genomic DNA isolated, and sgRNAs enriched by the respective selective pressure were identified via next-generation sequencing by comparison to respective untreated control samples. (**b**) Circos plot of haploid genetic screens in KBM7 cells with 250 ng/ml FASL (red) or 100 ng/ml TRAIL (blue). Each dot represents a mutagenized gene identified in the resistant cell population, the dot size corresponds to the number of independent insertions identified for each gene and the distance from center indicates the significance of enrichment compared to an unselected control data set. Hits with a *p*-value <10^-10^ are labelled by name. (**c**) MCA of KBM7 *SpCas9* cells transduced with a GFP marker (GFP^+^) and sgRNAs targeting either *SLC39A7* or *Renilla luciferase* (sg*Ren*) as control, against cells transduced with sg*Ren* and an mCherry marker (mCherry^+^). The cell populations were mixed in a 1:1 ratio, treated with TRAIL (100 ng/ml), FasL (20 ng/ml), or a combination of TNFα (10 ng/ml) and SMAC mimetic (0.5 µM), and analyzed after 14 days by flow cytometry. Data represent mean value ± s.d. of two independent experiments performed in duplicates, n.d. (not determined) indicates wells with no outgrowth. (**c**) Flow cytometry analysis for FAS surface expression in KBM7 *FADD^-^ SLC39A7*^-^ cells reconstituted with either SLC39A7 or GFP. KBM7 *FADD*^-^ sg*Ren* and empty KBM7 *FADD^-^ SLC39A7*^-^ cells serve as positive and negative control, respectively. Data shown are representative of two independent experiments. (**e**) Flow cytometry analysis for surface expression of the indicated markers in KBM7 *FADD^-^ SLC39A7*^-^ cells and KBM7 *FADD*^-^ sg*Ren* control. Data shown are representative of two independent experiments.

**Supplementary Figure 5:**
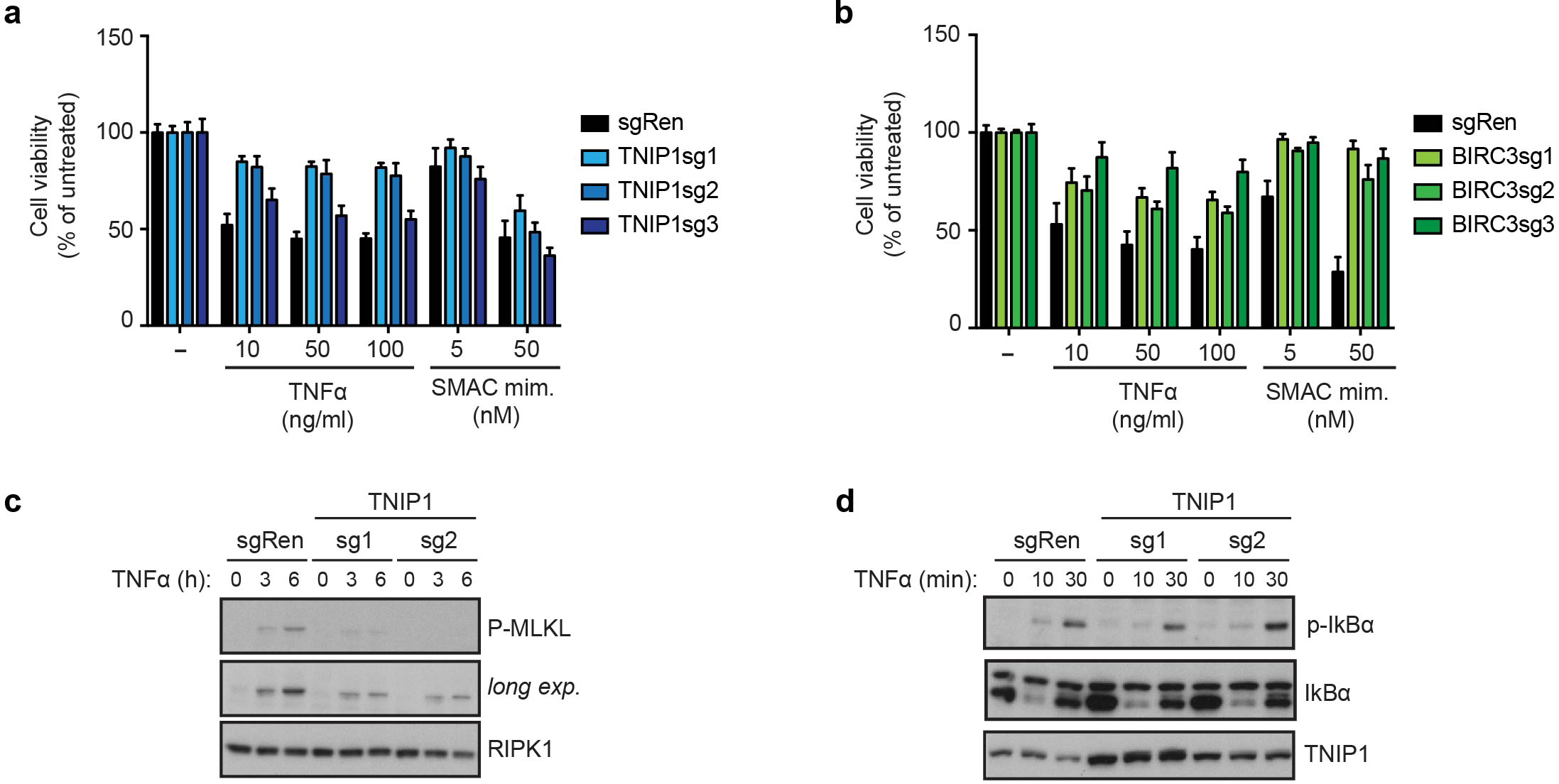
sgRNA–mediated transcriptional activation of TNIP1 or BIRC3 inhibits necroptosis. (**a-b**) Cell viability in KBM7 *FADD*^-^ SAM cells transduced with sgRNAs targeting Renilla control, TNIP1 (**a**) or BIRC3 (**b**). Cells were treated as indicated for 24h and viability was assessed using a luminescence-based readout for ATP (CellTiter Glo). Data represent mean value ± s.d. of two (**b**) or three (**a**) independent experiments performed in triplicates. (**c-d**) KBM7 *FADD*^-^ cells stably expressing sgRNAs targeting Renilla or TNIP1 were stimulated for the time indicated with TNFα (100ng/ml in **c**; 10 ng/ml in **d**). Cells were then lysed and subjected to immunoblotting with the indicated antibodies. Data shown are representative of two independent experiments.

**Supplementary Table 1: Haploid genetic screen results.** Tables listing the number of identified disruptive insertions per gene, inactivating insertions identified in other genes, total insertions in the control population, *p*-value, and adjusted *p*-value of enrichment for each screen.

**Supplementary Table 2: Proteomics data.** Tables listing the proteins identified by mass spectrometry in KBM7 *FADD^-^ SpCas9 SLC39A7*^-^ cells compared to sg*Ren* control, GO term enrichment analysis results in upregulated proteins (log2fold change >1) in KBM7 *FADD^-^ SpCas9 SLC39A7*^-^ cells compared to sg*Ren* control, and hits of GSEA using Hallmark gene sets on significantly up-or downregulated proteins (*p*-value <0.05) in KBM7 *FADD^-^ SpCas9 SLC39A7*^-^ cells compared to sg*Ren* control.

**Supplementary Table 3: sgRNA sequences used in this study.** Tables listing target genes and the respective sgRNA sequences used.

**Supplementary Table 4: SLC-focused KO genetic screen results.** Table listing the results of the enrichment analysis at sgRNA (DESeq2) and gene (GSEA) level for each screen.

**Supplementary Table 5: sgRNA counts of SLC-focused KO genetic screens.** Table listing sgRNA counts for each screen.

**Supplementary Table 6: Gain-of-function genetic screen results.** Table listing the results of the enrichment analysis at sgRNA (DESeq2) and gene (GSEA) level for each screen.

**Supplementary Table 7: sgRNA counts of the gain-of-function genetic screen.** Table listing sgRNA counts for each screen.

